# Signaling scaffold Shoc2 regulates lymphangiogenesis by suppressing mTORC1-mediated IFN responses

**DOI:** 10.1101/2025.03.26.645567

**Authors:** Patricia Wilson, Vishakha Vishwakarma, Rebecca Norcross, Kashmira Khaire, Van N. Pham, Brant M. Weinstein, Hyun Min Jung, Emilia Galperin

**Affiliations:** Department of Molecular and Cellular Biochemistry, University of Kentucky, Lexington, KY 40536; Division of Developmental Biology, Eunice Kennedy Shriver National Institute of Child Health and Human Development, NIH, Bethesda, MD 20814, USA; Clinical Operations R&D group, Zoetis, Kalamazoo, MI, 49001; Department of Molecular and Cellular Developmental Biology, University of California, Los Angeles, CA, 90095; Department of Pharmacology and Regenerative Medicine, University of Illinois at Chicago, Chicago, IL, 60612

## Abstract

An interplay of growth factors and signaling pathways governs the development and maintenance of the lymphatic vasculature, ensuring proper fluid homeostasis and immune function. Disruption of these regulatory mechanisms can lead to congenital lymphatic disorders and contribute to various pathological conditions. However, the mechanisms underlying the molecular regulation of these processes remain elusive. Here we reveal a critical and previously unappreciated role for the signaling scaffold protein Shoc2 in lymphangiogenesis. We demonstrate that loss of Shoc2 leads to nearly a complete loss of lymphatic vasculature in vivo and senescence of lymphatic endothelial cells in vitro. Mechanistically, Shoc2 is required for balancing signaling through the ERK1/2 pathway, and its loss results in increased mTORC1 signaling. This dysregulation impairs mitochondrial respiration and triggers an IRF/IFN-II response, ultimately leading to cellular senescence. Strikingly, expression of the Noonan Syndrome with Loose anagen Hair (NSLH)-causing Shoc2 variant S2G phenocopies the effects of Shoc2 loss. Together, these studies establish the critical role of Shoc2 in lymphangiogenesis and uncover a novel mechanistic link between Shoc2 signaling, mitochondrial function, innate immune response, and lymphatic development, with significant implications for Ras-pathway-related congenital disorders.

## INTRODUCTION

The lymphatic vasculature plays a multifaceted role that extends beyond the continuous collection of excess interstitial fluid and macromolecules and the transport of this fluid to the bloodstream. (1) A critical component of the immune system, the lymphatic system serves as a conduit for the trafficking of immune cells and is essential for both adaptive and innate immune responses. (2, 3) The intricate function of the lymphatic system, including its role in immune surveillance and response, relies heavily on the precise regulation of lymphatic endothelial cells (LECs). LECs typically reside in a quiescent state, characterized by minimal proliferative activity. This quiescent state, however, is a dynamic equilibrium actively maintained by a complex interplay of intracellular signaling pathways. Disrupting this delicate balance within LECs can have profound consequences. (3, 4) Aberrant activation of key signaling cascades, including the PI3K/AKT and Ras/ERK pathways, can drive LECs out of their quiescent state, inducing uncontrolled proliferation and lymphatic malformations that manifest as abnormal lymphatic vessel growth, localized swelling, or more severe conditions, which can significantly impact organ function. (5, 6)

The pivotal role of the Ras/ERK signaling pathway in LEC function is underscored by syndromes collectively known as RASopathies. (7) These spectrum disorders arise from mutations in the genes of the Ras/ERK pathway. More than 20% of RASopathy patients exhibit lymphatic dysfunction presenting in a variety of forms, including lymphedema, a chronic condition characterized by the accumulation of fluid in the interstitial spaces, and more severe conditions, such as chylothorax (fluid in the pleural cavity), hydropericardium (fluid around the heart), or chylous ascites (fluid in the abdomen). (8, 9) While lymphatic abnormalities in RASopathies have significant clinical implications, their causes are not fully understood, and it remains unclear why some lymphatic abnormalities may appear later in life and what triggers them.

Noonan Syndrome with loose anagen hair (NSLH) (OMIM #607721) is a RASopathy caused by hereditary variants in the *Shoc2* gene. (10, 11) NSLH patients manifest many typical RASopathy presentations, including lymphatic abnormalities. (11–15) Due to the rarity of NSLH, the literature describing the lymphatic presentation is limited, but several studies have reported that patients harboring the Shoc2 variant c. A4>G, p. S2G present with severe lymphatic dysfunction, including fetal hydrops, pleural effusions, and generalized edema resulting in death. (14, 16, 17)

Shoc2 is a non-redundant, evolutionarily conserved signaling scaffold protein and a key regulator of Ras/ERK1/2 pathway signal amplitude. Shoc2 tethers Ras and the catalytic subunit of protein phosphatase 1c (PP1c, also known as PP1CA) to dephosphorylate inhibitory phospho-S259 of RAF-1 and accelerate ERK1/2 signals. (18) Shoc2 other interacting partners include the proteins of the ubiquitin system HUWE1, VCP, USP7, PSMC5, and FBXW7 (11, 19–23), scaffold proteins SCRIB and Raptor, and signaling enzymes such as the p110α subunit of phosphatidylinositol 3-kinase (PI3K). (24) While the role of these interactions in distinct physiological processes or pathological conditions is unclear, Shoc2’s significance in embryonic development, morphogenesis, and the maintenance of various tissues in adult organisms is highlighted in vertebrate models. (25, 26) Shoc2 *null* mice and zebrafish CRISPR/Cas9 knockouts both develop systemic deficiencies resulting in early embryonic lethality. Interestingly, zebrafish CRISPR/Cas9 Shoc2 mutants develop unusual edemic swellings, while endothelial-specific Shoc2 knockout in mice causes sporadic hemorrhaging, subcutaneous edema, and fetal lung congestion, suggesting potential vascular deficits in these mutants. (25, 26)

Given the importance of Shoc2 in ERK1/2 signaling, we asked whether Shoc2 regulates lymphatic development and maintenance of lymphatic endothelial cells. Using an in vivo approach, we demonstrate that Shoc2 is vital for developmental lymphangiogenesis. Analysis of Shoc2 *null* zebrafish embryos shows a near-total lack of lymphatic vessels. The lack of lymphatic vessels can be rescued by endothelial-specific expression of wild-type Shoc2, demonstrating the endothelial cell-autonomous requirement of Shoc2 for lymphatic development. Our in vitro studies show that Shoc2 loss in primary human LECs impairs mTOR pathway activation, thereby disrupting mitochondrial function, activating nucleic acid sensing machinery, and the IFN/JAK1/STAT1-mediated innate immune response, ultimately leading to cell senescence. Re-expression of wild-type but not NSLH mutant Shoc2 in Shoc2-deficient LECs restores mitochondrial function and reverses innate immune responses. Taken together, this study uncovers the cell-type-specific and essential role of Shoc2 in orchestrating lymphangiogenesis and homeostasis of LECs and provides novel insights into the detailed molecular mechanisms potentially underpinning lymphatic dysfunction in NSLH.

## RESULTS

### Developmental abnormalities in Shoc2 mutant fish

To understand the role of Shoc2-mediated signaling during lymphatic development, we utilized a stable zebrafish line carrying a germline mutation in the *shoc2* gene. This line, ZDB-ALT-131217-17590, also known as sa24200 (obtained from ZIRC), carries a c.1546G>A mutation that disrupts normal splicing by inserting a 31-bp cryptic exon into the *Shoc2* mRNA (from here, the mutant allele is referred to as *shoc2^SA^*). The c.1546G>A mutation results in a premature stop codon (p. Val345*) after exon 5 (**Figure S1A-C**). We confirmed this aberrant splicing by sequencing cDNA isolated from 6 days post-fertilization (dpf) *shoc2^SA/SA^* larvae. Quantitative PCR revealed a significant reduction in *Shoc2* mRNA levels in *shoc2^SA/SA^*larvae compared to controls (**Figure S1D**), and immunoblotting confirmed the absence of Shoc2 protein in *shoc2^SA/SA^* larvae (**Figure S1E**). Heterozygous *shoc2^+/SA^* alleles did not show a decrease in Shoc2 protein levels. These results indicate that *shoc2^SA/SA^* is a loss-of-function mutation that causes nonsense-mediated *Shoc2* mRNA decay. *shoc2^SA/SA^* mutants were outcrossed to wild-type fish for at least three generations before further study.

Heterozygous *shoc2^+/SA^* carriers were viable and fertile with no apparent abnormalities. However, homozygous *shoc2^SA/SA^* larvae displayed severe developmental defects. At 5 dpf, nearly 80% of *shoc2^SA/SA^* larvae exhibited edema throughout the whole body, most notably in the heart cavity, trunk, yolk sac, yolk extension, and around the eyes. They also had underinflated swim bladders and were lethargic. These phenotypes closely resemble those observed in our previously described *shoc2^Δ22^* mutant line (**Figure 1A, B**, and **Figure S1F**). (25) A complementation test using two mutant alleles, *shoc2^Δ22^* and *shoc2^SA/SA^*, revealed that a severe edema phenotype can only be found in the *shoc2^Δ22/SA^* larvae, confirming that the loss of Shoc2 specifically drives the edema phenotype. Survival studies showed that *shoc2^SA/SA^* larvae began to die at 6 dpf, with none surviving past 11 dpf (**Figure S1G**). These data demonstrate that the c.1546G>A substitution in *shoc2* severely impairs zebrafish embryonic development.

**Figure 1.**
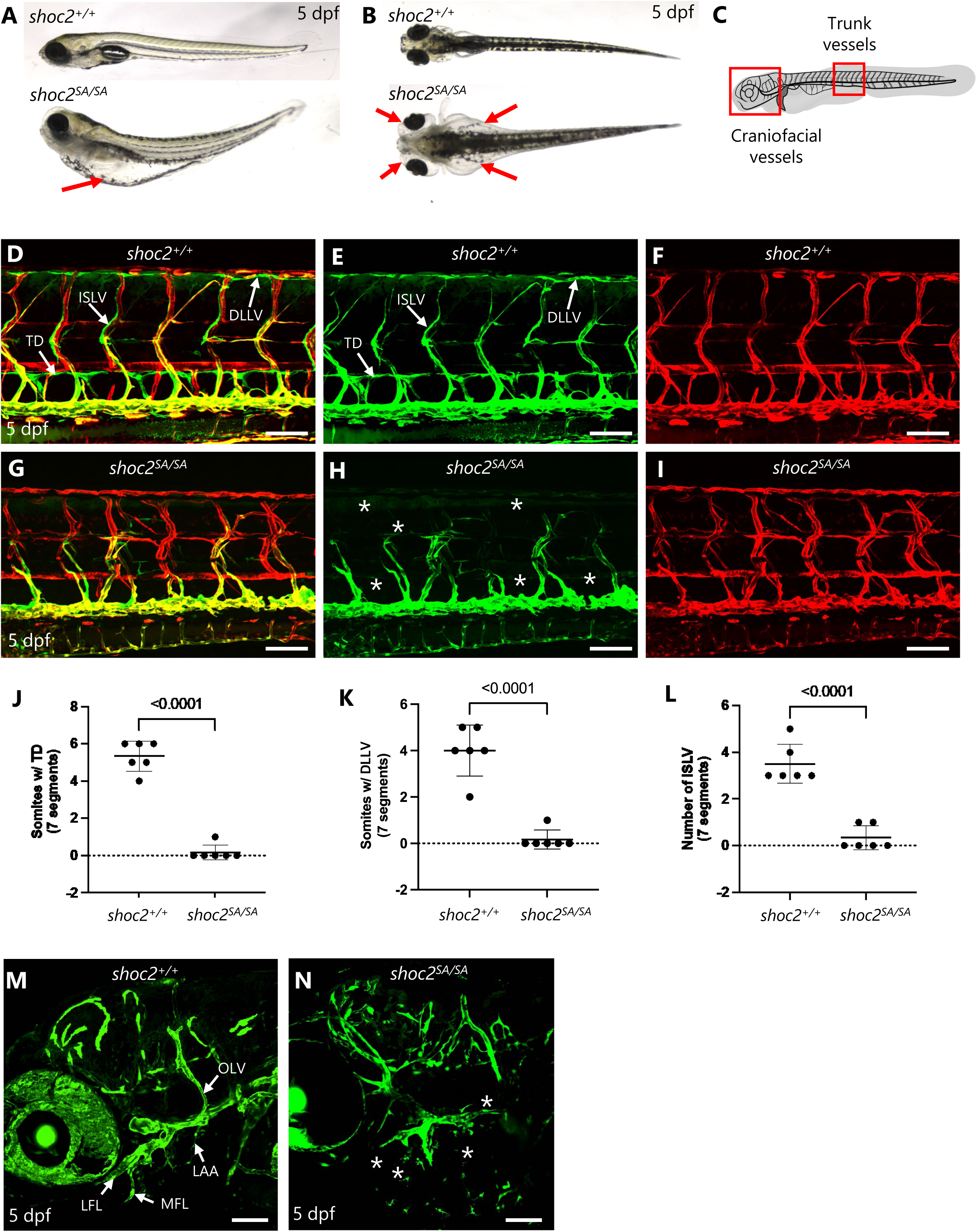
Loss of lymphatics in shoc2 mutant zebrafish. **A.** Brightfield images of a lateral view of the entire zebrafish larvae of wild-type sibling (top panel) and *shoc2^SA/SA^* mutant (bottom panel). The red arrow is pointing to an edematous site. **B.** Brightfield images of a dorsal view of the entire zebrafish larvae of wild-type sibling (top panel) and *shoc2^SA/SA^* mutant (bottom panel). Red arrows are pointing to edematous sites. **C.** Schematic of areas of confocal imaging highlighting the trunk and craniofacial vessels. **D.** Confocal image of trunk vessels of wild-type sibling *Tg(mrc1a:eGFP)^y251^;Tg(kdrl:mCherry)^y171^*zebrafish larvae at 5 dpf. Scale bar: 100 μm. TD, thoracic duct; ISLV, intersegmental lymphatic vessel; DLLV, dorsal longitudinal lymphatic vessel. **E.** Confocal image of trunk vessels of a wild-type sibling *Tg(mrc1a:eGFP)^y251^* showing a separated green channel from panel D. Scale bar: 100 μm. **F.** Confocal image of trunk vessels of a wild-type sibling *Tg(kdrl:mCherry)^y171^* showing a separated red channel from panel D. Scale bar: 100 μm. **G.** Confocal image of trunk vessels of a *shoc2^SA/SA^* mutant *Tg(mrc1a:eGFP)^y251^;Tg(kdrl:mCherry)^y171^* zebrafish larvae at 5 dpf. Asterisks indicate the absence of lymphatic vessels. Scale bar: 100 μm. **H.** Confocal image of trunk vessels of a *shoc2^SA/SA^* mutant *Tg(mrc1a:eGFP)^y251^* showing separated green channel from panel D. Scale bar: 100 μm. **I.** Confocal image of trunk vessels of a *shoc2^SA/SA^* mutant *Tg(kdrl:mCherry)^y171^* showing separated red channel from panel D. Scale bar: 100 μm. **J.** Quantitation of the number of somites with TD segments in wild-type or *shoc2^SA/SA^* mutants at 5 dpf (n=6 larvae in each group). **K.** Quantitation of the number of somites with DLLV segments in wild-type or *shoc2^SA/SA^* mutants at 5 dpf (n=6 larvae in each group). **L.** Quantitation of the number of somites with ISLV in wild-type or *shoc2^SA/SA^* mutants at 5 dpf (n=6 larvae in each group). **M.** Confocal image of the craniofacial lymphatics of a wild-type sibling at 5 dpf. LFL, lateral facial lymphatics, MFL, medial facial lymphatics, OLV, otolithic lymphatic vessel, LAA, lymphatic branchial arches. Scale bar: 100 μm. **N.** Confocal image of the craniofacial lymphatics of a *shoc2^SA/SA^* mutant at 5 dpf. Asterisks indicate the absence of lymphatic vessels. Scale bar: 100 μm.

Morphological analysis of *shoc2^SA/SA^* mutants revealed craniofacial abnormalities, including a prominent defect in Meckel’s cartilage (curved downwards and shorter than in wild-type larvae) and hypoplastic closure of Meckel’s cartilage. Chondrocyte stacking within the cartilaginous elements of *shoc2^SA/SA^* was also disorganized (**Figure S2A**). Alizarin Red S staining showed significantly reduced calcification of craniofacial bones in *shoc2^SA/SA^* compared to wild-type controls (**Figure S2B**). *shoc2^SA/SA^* mutants also displayed abnormal pigmentation, including a loss of iridophores (**Figure S2C, D**) and irregular melanocyte patterning (**Figure S2E**). Collectively, these data demonstrate that the *shoc2* c.1546G>A variant phenocopies the *shoc2^Δ22^* mutant (25), making it a suitable model for further investigation of Shoc2’s role in zebrafish development.

### Defective lymphatic development in Shoc2 mutant fish

Zebrafish are widely used as a vertebrate model for studying lymphatic development due to their optical clarity, external development, high conservation of the lymphatic vascular system, and the availability of transgenic lines expressing fluorescent reporters in LECs that enable *in vivo* monitoring of lymphatic development. (27, 28). To investigate the role of Shoc2 in lymphatic development and its relation to the edematous phenotype, we crossed the *shoc2^SA/SA^* mutants with *Tg(mrc1a:eGFP)^y251^;Tg(kdrl:mCherry)^y171^*double transgenic zebrafish. This line marks lymphatics, arteries, and veins with green, red, or both fluorescent proteins, respectively. (28) We focused on two major lymphatic vascular beds, the trunk and craniofacial lymphatic networks (**Figure 1C**).

At 5 dpf, wild-type larvae exhibited normal lymphatic development, including the thoracic duct (TD), intersegmental lymphatic vessels (ISLV), and dorsal longitudinal lymphatic vessels (DLLV) (**Figure 1D-F**). In contrast, *shoc2^SA/SA^* mutants showed a marked loss of lymphatic vessels, including the TD, ISLV, and DLLV (**Figure 1G, H**), while blood vessel development remained largely unaffected, indicating that the defects are specific to the lymphatic endothelial cells (**Figure 1I**). Quantification of TD, ISLV, and DLLV formation in wild-type siblings and *shoc2^SA/SA^* mutants revealed a nearly complete loss of all three sets of trunk lymphatic vessels in Shoc2-deficient animals (**Figure 1J-L**). Craniofacial lymphatic vessels were also strongly reduced in the mutants (**Figure 1M, N**). These findings indicate that lymphatic development is disrupted in Shoc2-deficient zebrafish.

### Loss of lymphatic progenitors in Shoc2 mutant fish

Lymphatic network formation in the zebrafish trunk begins with secondary sprouts that emerge from the posterior cardinal vein around 1.5 dpf. (29, 30) By 3 dpf, these sprouts contribute to the formation of the parachordal line (PL, or parachordal angioblasts), a transient progenitor structure that gives rise to the trunk lymphatic vessels (**Figure 2A**). Using time-lapse imaging of *Tg(mrc1a:eGFP)^y251^;Tg(kdrl:mCherry)^y171^* transgenic fish, we analyzed the appearance of secondary sprouts from 32 hours post-fertilization (hpf) to 50 hpf in wild-type siblings (**Figure 2B-D**) and Shoc2 mutants (**Figure 2E-G**). In wild-type siblings, secondary sprouts emerged from the posterior cardinal vein and migrated along the intersegmental blood vessels (**Figure 2B-D** and **Video S1**). In contrast, *shoc2^SA/SA^* mutants produced only small abortive secondary sprouts up to 50 hpf (**Figure 9E-G** and **Video S2**). By 3 dpf, wild-type siblings formed PLs in the horizontal myoseptum region (**Figure 9H, I**), while *shoc2^SA/SA^* mutants failed to form PLs (**Figure 2J, K**, asterisks). Quantification of PLs confirmed a near-total loss of PL lymphatic progenitors in *shoc2^SA/SA^* mutants compared to wild-type siblings (**Figure 2L**).

**Figure 2.**
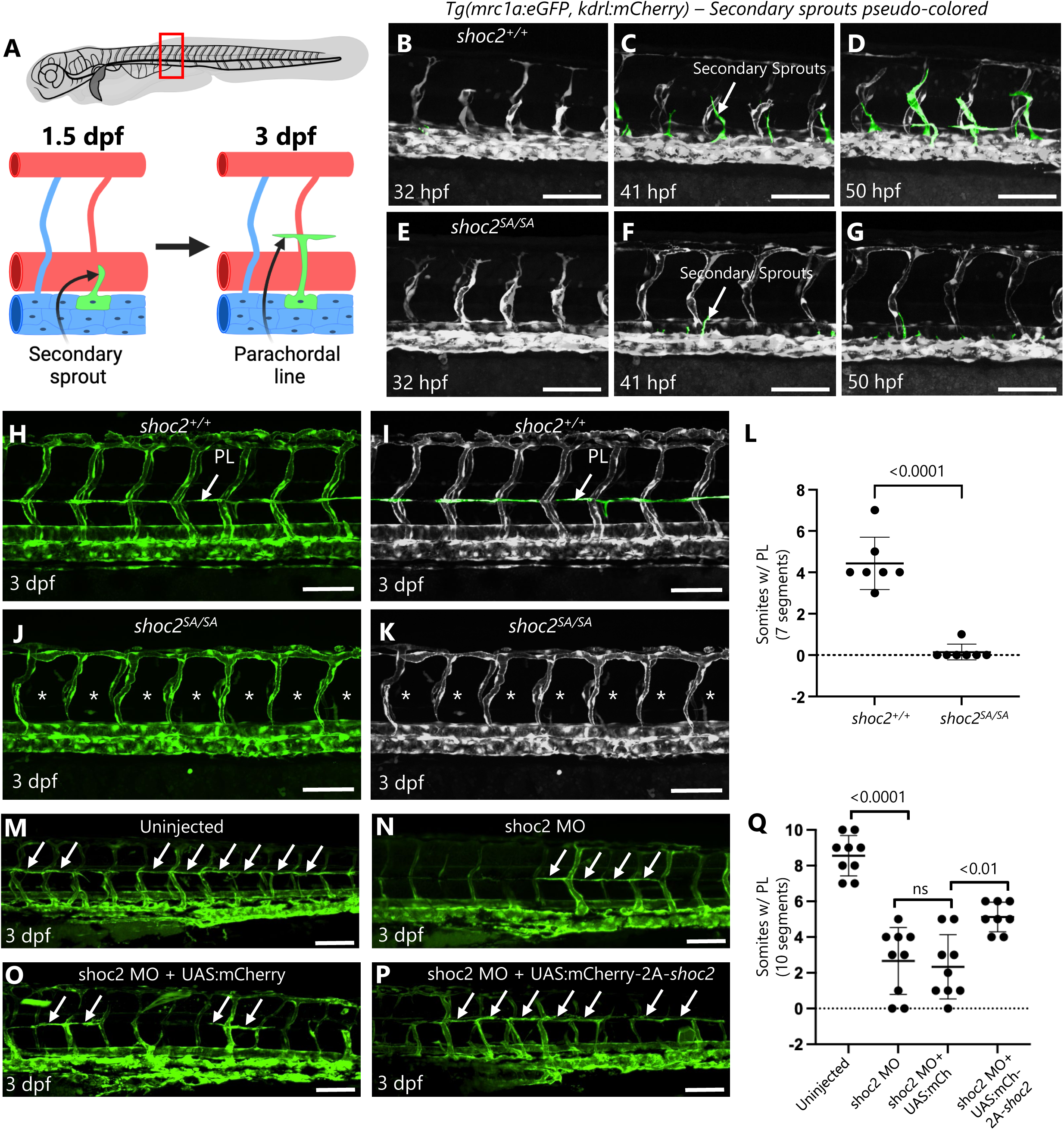
Defective lymphatic progenitors in shoc2 mutant zebrafish. **A.** A schematic showing the region of interest where lymphatic vessels originate. Secondary sprouts emerge from a posterior cardinal vein (PCV) and migrate dorsally around 1.5 dpf, giving rise to the parachordal lines (PL) at 3 dpf. These PLs are lymphatic progenitors that migrate and proliferate to form the trunk lymphatic vessels. **B.** Confocal image of trunk vessels of a wild-type sibling *Tg(mrc1a:eGFP)^y251^;Tg(kdrl:mCherry)^y171^* zebrafish larvae at 32 hpf. The image is converted to a grey scale. Scale bar: 100 μm. **C.** Confocal image of trunk vessels of a wild-type sibling *Tg(mrc1a:eGFP)^y251^;Tg(kdrl:mCherry)^y171^*zebrafish larvae at 41 hpf. The image is converted to a grey scale, and secondary sprouts are pseudocolored in green. Scale bar: 100 μm. **D.** Confocal image of trunk vessels of a wild-type sibling *Tg(mrc1a:eGFP)^y251^;Tg(kdrl:mCherry)^y171^*zebrafish larvae at 50 hpf. The image is converted to a grey scale, and secondary sprouts are pseudocolored in green. Scale bar: 100 μm. **E.** Confocal image of trunk vessels of a *shoc2^SA/SA^* mutant *Tg(mrc1a:eGFP)^y251^;Tg(kdrl:mCherry)^y171^* zebrafish larvae at 32 hpf. The image is converted to a grey scale. Scale bar: 100 μm. **F.** Confocal image of trunk vessels of a *shoc2^SA/SA^* mutant *Tg(mrc1a:eGFP)^y251^;Tg(kdrl:mCherry)^y171^*zebrafish larvae at 41 hpf. The image is converted to a greyscale, and secondary sprouts are pseudocolored in green. Scale bar: 100 μm. **G.** Confocal image of trunk vessels of a *shoc2^SA/SA^* mutant *Tg(mrc1a:eGFP)^y251^;Tg(kdrl:mCherry)^y171^*zebrafish larvae at 50 hpf. The image is converted to a greyscale, and secondary sprouts are pseudocolored in green. Scale bar: 100 μm. **H.** Confocal image of trunk vessels of a wild-type sibling *Tg(mrc1a:eGFP)^y251^;Tg(kdrl:mCherry)^y171^* zebrafish larvae at 3 dpf. PL, parachordal line. Scale bar: 100 μm. **I.** Greyscale image of a wild-type sibling *Tg(mrc1a:eGFP)^y251^;Tg(kdrl:mCherry)^y171^* in panel H. PLs are pseudocolored in green. Scale bar: 100 μm. **J.** Confocal image of trunk vessels of a *shoc2^SA/SA^* mutant *Tg(mrc1a:eGFP)^y251^;Tg(kdrl:mCherry)^y171^* zebrafish larvae at 3 dpf. Asterisks indicate the absence of PLs. Scale bar: 100 μm. **K.** Greyscale image of a *shoc2^SA/SA^* mutant *Tg(mrc1a:eGFP)^y251^;Tg(kdrl:mCherry)^y171^* in panel J. Asterisks indicate the absence of PLs. Scale bar: 100 μm. **L.** Quantification of the number of somites containing PL segments in wild-type or *shoc2^SA/SA^* mutants at 3 dpf. Seven somites per larvae were counted (n=7 larvae in each group). **M.** Confocal image of trunk vessels of an uninjected *Tg(mrc1a:kalTA4, UAS:eGFP)* zebrafish larvae at 3 dpf. The arrows indicate PLs. Scale bar: 100 μm. **N.** Confocal image of trunk vessels of a *Tg(mrc1a:kalTA4, UAS:eGFP)* shoc2 MO-injected zebrafish larvae at 3 dpf. The arrows indicate PLs. Scale bar: 100 μm. **O.** Confocal image of trunk vessels of a *Tg(mrc1a:kalTA4, UAS:eGFP)* shoc2 MO and UAS:mCherry co-injected zebrafish larvae at 3 dpf. The arrows indicate PLs. Scale bar: 100 μm. **P.** Confocal image of trunk vessels of a *Tg(mrc1a:kalTA4, UAS:eGFP)* shoc2 MO and UAS:mCherry-2A-shoc2 co-injected zebrafish larvae at 3 dpf. The arrows indicate PLs. Scale bar: 100 μm. **Q.** Quantification of the number of somites containing PL segments in panels M-P. Ten somites per larvae were counted. Data points indicate the number of larvae in each group.

We also examined the development of the dorsal aorta-derived primary sprouts that generate intersegmental blood vessels by performing time-lapse imaging of *Tg(fli1:eGFP)^y1^* transgenic zebrafish from 24 to 36 hpf in both wild-type siblings and *shoc2^SA/SA^*mutants (**Video S3, 4**). Primary sprouts formed normally in both genotypes, suggesting that Shoc2 loss-of-function primarily affects secondary sprout formation but not primary sprouts (**Figure S3)**. These findings indicate that the lymphatic defects in *shoc2^SA/SA^* mutants are due to the lack of secondary sprout formation and the subsequent failure to produce lymphatic progenitor cells during early larval development.

### Endothelial expression of wild-type Shoc2 rescues lymphatic development in Shoc2 mutants

To examine whether endothelial Shoc2 function is sufficient to support lymphatic development, we drove mosaic transgenic expression of wild-type Shoc2, specifically in venous and lymphatic endothelial cells. We generated *Tol2(uas:mcherry-2A-shoc2)* and control *Tol2(uas:mcherry)* transgenes and injected them separately into single-cell *Tg(mrc1a:KalTA4; UAS:eGFP)* transgenic animals together with a morpholino targeting the translation start site of endogenous but not transgene-driven Shoc2. The animals were raised and examined for PL formation at 3 dpf. As previously reported, morpholino injection phenocopied Shoc2 mutants, resulting in a dramatic loss of PL formation compared to uninjected siblings (**Figure 2M, N, and Q**). Co-injection of the control *Tol2(uas:mcherry)* transgene did not mitigate the effects of the morpholino on lymphatic development, whereas co-injection of the Shoc2-expressing *Tol2(uas:mcherry-2A-shoc2)* transgene strongly rescued parachordal line formation in Shoc2 morphants (**Figure 2P**). These results show that Shoc2 function is required autonomously within endothelial cells to permit lymphatic development to proceed.

### Shoc2 controls the amplitude of VEGF-mediated ERK1/2 signals

Recognizing the challenge posed by the near-complete loss of lymphatic vessels in zebrafish Shoc2 *null* larvae for studying LEC homeostasis, we employed human primary lymphatic endothelial cells (HDLEC) as a tractable *in vitro* system to investigate Shoc2 function in these cells. First, we determined that Shoc2 regulates VEGF-initiated activation of the ERK1/2 pathway in HDLEC and human umbilical vein endothelial cells (HUVEC). Shoc2 shRNA depletion (KD) significantly decreased ERK1/2 phosphorylation following stimulation with VEGF-C (HDLEC) or VEGF-A (HUVEC) in both cell types. Conversely, ectopic expression of Shoc2 tagged with tagRFP (SE) rescued ERK1/2 phosphorylation, indicating that Shoc2 is important for VEGF-mediated ERK1/2 signaling in endothelial cells (**Figures 3A**-**D**). Interestingly, we observed notable changes in the morphology of Shoc2-depleted HDLEC cultured for more than 6 days. To characterize this phenotype, we examined senescence-associated β-galactosidase (SA-β-gal) activity. As shown in **Figure 3E**, almost the entire population of Shoc2-depleted HDLEC was positive for SA-β-gal. Moreover, cell size distribution analysis indicated that Shoc2-depleted cells have larger cell volumes (**Figure 3F**). Shoc2-depleted HDLEC also had increased levels of p53 and phospho-p53, as well as increased levels of senescence-associated cell cycle arrest markers p21^Cip1^ (**Figure 3G**). Together, these data indicate a senescence phenotype in Shoc2-depleted HDLEC. Yet, Shoc2 loss had a minimal impact on HUVEC (**Figure 3G**), suggesting a stringent requirement of Shoc2 for HDLEC proliferation and growth, and implying that the downstream targets of Shoc2-mediated signals potentially involve distinct effector proteins or signaling pathways in the two endothelial cell types.

**Figure 3.**
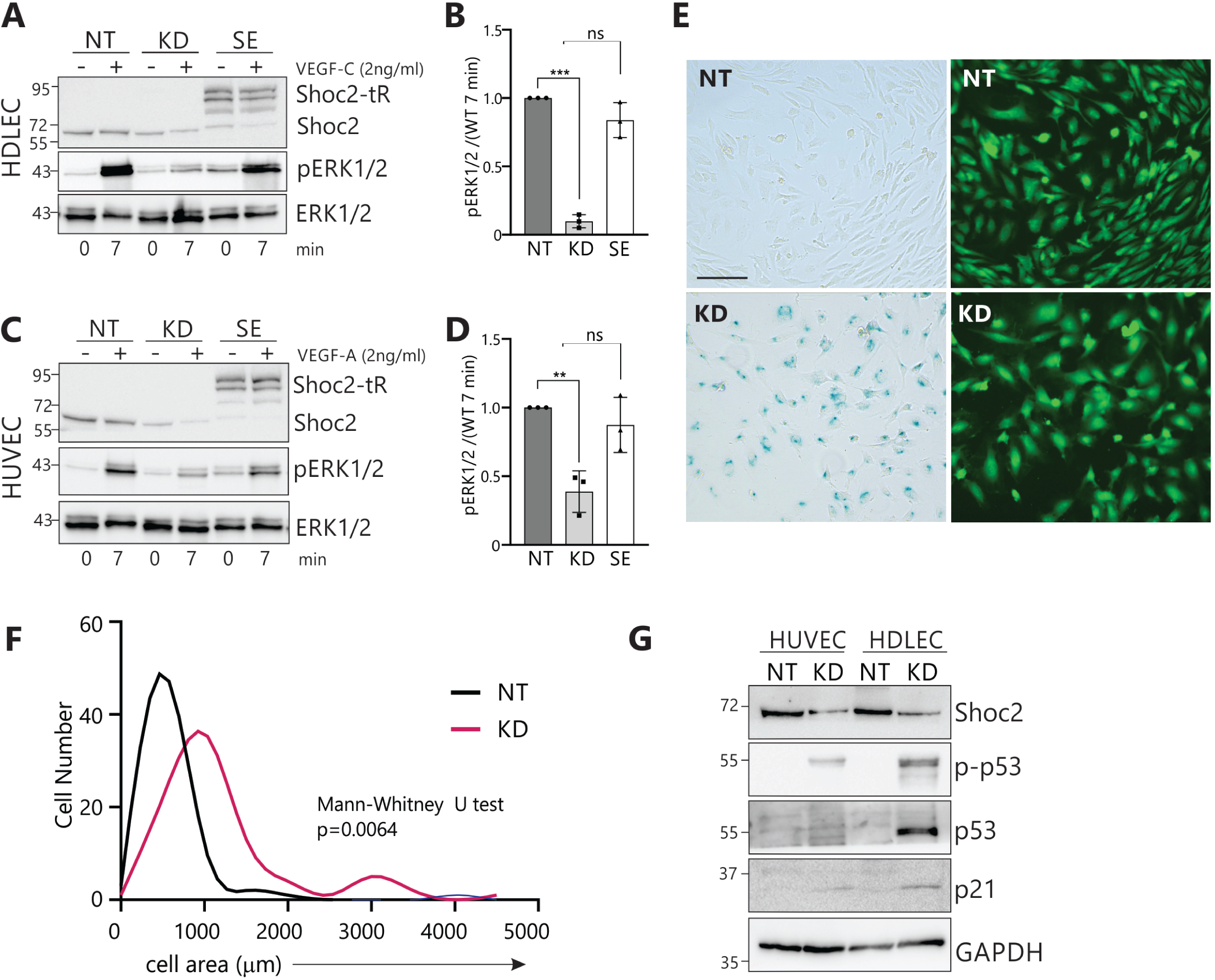
Shoc2-dependent activation of the ERK1/2 pathway in endothelial cells. **A.** Western blot of whole cell lysates extracted from human dermal lymphatic endothelial cells (HDLEC) expressing lentivirally supplied nontargeting shRNA (NT), Shoc2 shRNA (KD) or Shoc2 shRNA together with Shoc2-tagRFP (SE), serum-starved for 16 hr and then stimulated or not with VEGF-C (2 ng/ml) for 7 min at 37°C. Cell lysates were probed using anti-pERK1/2, -ERK1/2, and - Shoc2 antibodies. **B.** Levels of pERK1/2 in cells treated with 2 ng/ml VEGFa at 7 min, normalized to the total amount of ERK1/2 and compared to pERK1/2 levels in parental HDLEC cells expressing nontargeting (NT) treated with 2 ng/ml VEGFa (set at 1.0). Values are shown in arbitrary units ± S.D. (*n*=3) (\**p =* 0.01 to 0.05, \*\**p* = 0.001 to 0.01, \*\*\**p =* 0.0001 to 0.001, \*\*\*\**p* < 0.0001, ≥ 0.05-non-significant, ns, one-way ANOVA Dunnett’s multiple comparison test). **C.** Western blot of whole cell lysates extracted from human umbilical vein endothelial cells (HUVEC) expressing lentivirally supplied nontargeting shRNA (NT), Shoc2 shRNA (KD) or Shoc2 shRNA together with Shoc2-tagRFP (SE), serum-starved for 16 hrs and then stimulated or not with VEGF-A (2 ng/ml) for 7 min at 37°C. Cell lysates were probed using anti-pERK1/2, -ERK1/2, and -Shoc2 antibodies. **D.** Levels of pERK1/2 in cells treated with 2 ng/ml VEGFa at 7 min, normalized to the total amount of ERK1/2 and compared to pERK1/2 levels in parental HUVEC cells expressing nontargeting (NT), treated with 2 ng/ml VEGFa (set at 1.0). Values are shown in arbitrary units ± S.D. (*n*=3) (\**p =* 0.01 to 0.05, \*\**p* = 0.001 to 0.01, \*\*\**p =* 0.0001 to 0.001, \*\*\*\**p* < 0.0001, ≥ 0.05-non-significant, ns, one-way ANOVA Dunnett’s multiple comparison test). **E.** HDLEC expressing lentivirally-supplied nontargeting shRNA (NT) or Shoc2 shRNA (KD) were stained to examine β-Galactosidase activity (left panels). GFP, expressed independently by the lentivirus, served as a reporter to monitor delivery efficiency (right panels). Scale bar – 150μm. **F.** Size distribution of HDLEC expressing lentivirally-supplied nontargeting shRNA (NT) or Shoc2 shRNA (KD), 6 days after transfection, as measured by surface area. **G.** Western blot of whole cell lysates extracted from HUVECs and HDLECs expressing lentivirally supplied nontargeting shRNA (NT) or Shoc2 shRNA (KD) for 6 days. Cell lysates were probed using anti-Shoc2, -p53, -53, and -p21 antibodies. The results in each panel are representative of those from three independent experiments.

### Transcription changes in Shoc2-depleted HDLEC

To explore the underlying mechanisms of Shoc2 depletion-induced cellular senescence, we performed RNA sequencing of HDLECs expressing nontargeting siRNA (NT, control), cells depleted of Shoc2 using shRNA (KD), or cells expressing both Shoc2 shRNA and siRNA-insensitive Shoc2-tagRFP (SE, rescue) (**Figure 4**). Three days post-transfection, when Shoc2 levels were significantly reduced (validated by western blot, **Figure S4A**), mRNA was isolated from each group for transcriptome profiling. Thorough DESeq2 analysis identified 316 differentially expressed genes (DEGs, fold change ≥ 0.5) across pairwise comparisons of Shoc2 KD and SE cells (Venn diagram, **Figure S4B**). We focused on genes differentially expressed between control (NT) and Shoc2-depleted (KD) cells.

**Figure 4.**
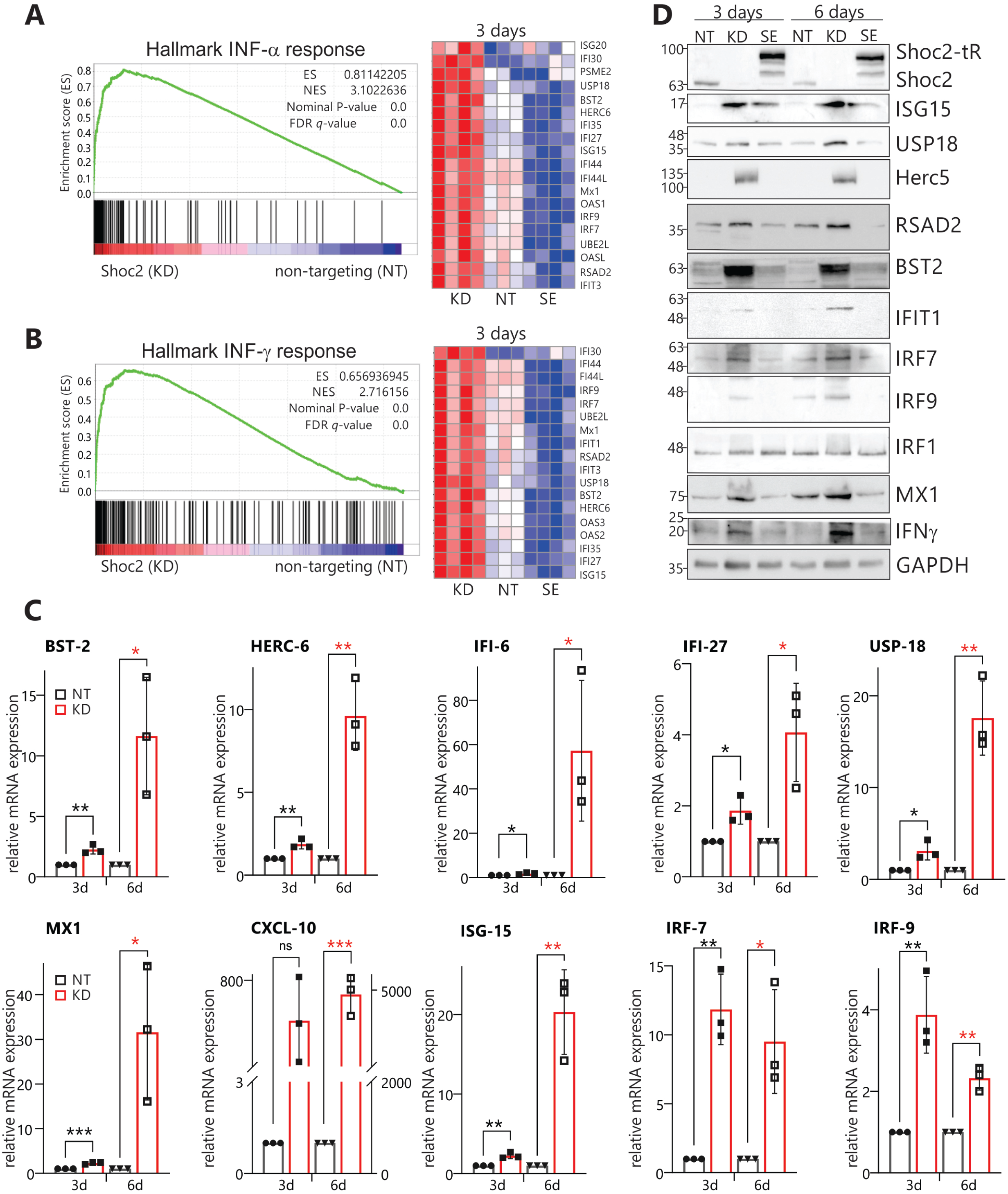
The Shoc2-dependent interferon response in HDLEC. **A-B.** Gene set enrichment analysis (GSEA) is presented as a score plot with the normalized enrichment score (NES) showing upregulation of interferon (IFN) response signature in Shoc2-depleted HDLEC (Shoc2 KD, KD) compared with control (nontargeting, NT) or HDLEC depleted of endogenous Shoc2 and expressing Shoc2-tagRFP (SE). The total height of the curve indicates the extent of enrichment (ES), with the normalized enrichment score (NES), the false discovery rate (FDR), and the p-value. Source data are provided as a source data file. Heatmap: the top elected correlated genes for each biological triplicate are presented in the corresponding heat map. Red and blue indicate increased and decreased expression for all heatmaps, respectively. **C.** The relative GAPDH-normalized expression of selected genes was measured by RT-qPCR. Data represent the mean and standard error of three independent experiments’ mean (SEM) values. P-values were calculated using a *t*-test (\**p =* 0.01 to 0.05, \*\**p* = 0.001 to 0.01, \*\*\**p =* 0.0001 to 0.001, \*\*\*\**p* < 0.0001, ≥ 0.05-non-significant, ns). Black bars represent control (NT), and red bars represent Shoc2-depleted cells (KD). **D.** Cell lysates of Shoc2-depleted (KD), control (NT), or Shoc2-tagRFP expressing (SE) HDLEC were analyzed using western blot 3 days and 6 days post-transfection to determine the expression levels of the indicated proteins. The results in each panel are representative of at least three independent experiments.

Gene Set Enrichment Analysis (GSEA) using the “Hallmark” database uncovered negative enrichment of gene sets associated with “Cholesterol Homeostasis,” “Fatty acid metabolism,” “Peroxisome,” and “MYC targets” (normalized enrichment scores (NES) of -2.28, -1.93, -1.87, and -1.80 respectively). Intriguingly, we observed significant upregulation of “Interferon-alpha response” and “Interferon-gamma response” genes in Shoc2-depleted cells (normalized enrichment scores (NES) of 2.72 and 3.10, respectively) (**Fig. 4A, B,** and **Figure S4C**). Furthermore, we found significant enrichment of gene sets related to “Negative regulation of viral processes,” “Antigen processing and presentation,” and “Defense response to virus” (normalized p-value = 0.00) using the “Biological Processes” database analysis (**Figure S5A** and **B**). Detailed analysis confirmed increased expression of key genes within the IFN response pathways (**Fig. 4A, B, Figure S5C, D**), including members of the Interferon-Induced Protein with Tetratricopeptide Repeats (IFIT) gene family (e.g., *IFIT1*, *IFIT3*, *IFI27*, *IFI6*, *IFIT30*, *IFIT35*), Interferon-Stimulated Genes (ISGs) such as *ISG15* and *ISG20*, and genes regulating ISGylation (e.g., *USP18* and *HERC6*). (31, 32) Many of these upregulated genes contain interferon-sensitive response element (ISRE) *cis*-regulatory sites (e.g., *IFIT3*, *MX1*, *ISG15*) or both ISREs and gamma-activated sequences (GAS) (e.g., *BST2*, *IRF7*, and *OAS1*). (33–35)

qRT-PCR analysis of mRNA isolated from Shoc2-depleted HDLEC 3 and 6 days after transfection corroborated the RNA-seq data (**Figure 4C**) and demonstrated that Shoc2 loss also impacted the expression of lymphatic vascular markers *PROX1* and *LYVE1* (**Figure S6**). Furthermore, the changes in expression of examined genes were significantly more pronounced at 6 days post-transfection, suggesting an accumulating IFN response upon Shoc2 depletion. Immunoblot analysis confirmed increased expression of IFN response proteins in Shoc2-depleted HDLECs (**Figure 4D**). Shoc2-tagRFP expression rescued the IFN response, confirming the specificity of the Shoc2-depletion-induced gene expression changes. In Shoc2-depleted HDLEC, we also observed a significant increase in the protein levels of IRF7 and IRF9, transcription factors that regulate the production of interferons (IFNs), including IFN-γ. (36) These results indicate that Shoc2 loss in HDLECs induces active transcriptional changes associated with innate immune response.

### Shoc2 loss in HDLEC leads to activation of the JAK1/STAT1 intracellular signaling pathway

Type I and II interferons (IFNs) primarily signal through the Janus-Activated Kinase (JAK)-Signal Transducer and Activator of Transcription (STAT) pathway. (37) As illustrated in **Figure 5A**, this pathway involves JAK activation at the plasma membrane (PM), followed by rapid phosphorylation and nuclear translocation of STAT proteins. These phosphorylated STAT dimers then bind to interferon-stimulated response elements (ISREs) or gamma-activated sequences (GAS) on DNA, initiating the transcription of interferon-stimulated genes (ISGs). Thus, we investigated whether Shoc2 loss in HDLECs activates the JAK1/STAT1 pathway. Shoc2 depletion significantly increased the phosphorylation of JAK1 (Y1034/1035) and STAT1 (Y701), while expressing Shoc2-tagRFP abrogated this phosphorylation (**Figure 5B**). These findings confirm that in Shoc2-depleted HDLECs, the IFN response coincides with JAK1/STAT1 pathway activation.

**Figure 5.**
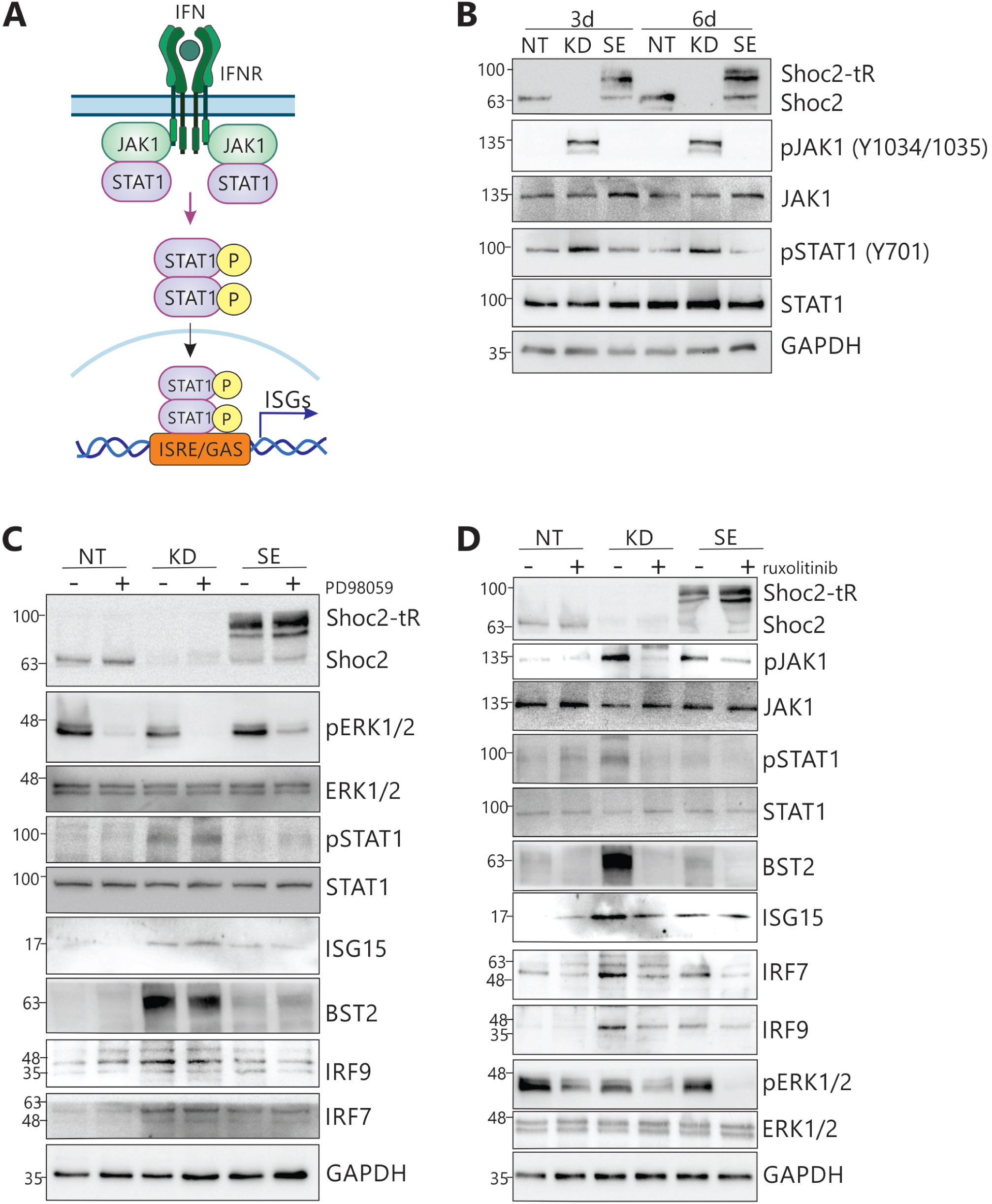
The ERK1/2 pathway-independent activation of the JAK1/STAT1 pathway in Shoc2-depleted HDLEC. **A.** Schematic representation of the major signaling pathway activated by IFNs - the Janus kinase/signal transducer and activator of transcription (JAK1/STAT1) pathway. Activation of the JAK1/STAT1 pathway at the plasma membrane and rapid phosphorylation and nuclear translocation of STAT1 leads to the expression of IFN-stimulated genes (ISGs) containing ISREs or GAS DNA cis-elements. **B.** Western blot of whole cell lysates extracted from human dermal lymphatic endothelial cells (HDLEC) expressing lentivirally supplied nontargeting shRNA (NT), Shoc2 shRNA (KD) or Shoc2 shRNA together with Shoc2-tagRFP (SE) for 3 and 6 days. Cell lysates were probed using antibodies to the indicated proteins. **C.** Western blot of whole cell lysates extracted from human dermal lymphatic endothelial cells (HDLEC) expressing lentivirally supplied nontargeting shRNA (NT), Shoc2 shRNA (KD) or Shoc2 shRNA together with Shoc2-tagRFP (SE) for 6 days. Cells were treated with 10µM MEK inhibitor PD50042 for 24 hours before western blot. Cell lysates were probed using antibodies to the indicated proteins. **D.** Western blot of whole cell lysates extracted from human dermal lymphatic endothelial cells (HDLEC) expressing lentivirally supplied nontargeting shRNA (NT), Shoc2 shRNA (KD) or Shoc2 shRNA together with Shoc2-tagRFP (SE) for 6 days. Cells were treated with 5µM JAK1 inhibitor ruxolitinib for 24 hours before western blot. Cell lysates were probed using antibodies to the indicated proteins. The results in each panel are representative of at least three independent experiments.

Earlier studies reported that ERK1/2 may regulate the expression of CCAAT/enhancer-binding protein-β (C/EBP-β), a transcription factor that governs the expression of various ISGs. (38, 39) We, therefore, tested whether inhibiting ERK1/2 activation would mimic the IFN response observed in the Shoc2-depleted HDLECs. **Figure 5C** shows that treatment of Shoc2-depleted, Shoc2-tagRFP-expressing, and control HDLECs with the MEK1/2 inhibitor PD98059 reduced overall levels of ERK1/2 phosphorylation but did not affect either STAT1 phosphorylation or the expression of IFN response proteins (i.e., BST2, ISG15, IRF7, and IRF9). Conversely, the JAK1 inhibitor ruxolitinib effectively eliminated the IFN response and activation of the JAK1/STAT1 pathway in Shoc2-depleted HDLEC (**Figure 5D**). Reduced ERK1/2 phosphorylation observed in cells treated with ruxolitinib is likely a downstream effect of JAK1/STAT1 pathway inhibition, as suggested in (40). These results demonstrate that IFN response and JAK-STAT pathway activation in HDLEC depleted of Shoc2 are independent of ERK1/2 pathway activity.

Next, we investigated whether Shoc2 loss induces a similar IFN/JAK1/STAT1 response in human umbilical vein endothelial cells (HUVECs). We also examined how the common Shoc2 NSLH-causing variant S2G impacts IFN/JAK1/STAT1 response (**Figure S7**). As expected, Shoc2 loss in HDLECs led to elevated levels of ISGs (i.e., ISG15, MX1, BST2, RSAD2), the transcription factors IRF7 and IRF9, as well as IFN-γ and STAT1 phosphorylation. Notably, in HDLEC depleted of Shoc2 and expressing the Shoc2 S2G variant (S2G), the IFN response and activation of the JAK1/STAT1 pathway were comparable to those of the Shoc2-depleted HDLEC (KD) (**Figure S7A**). Conversely, the IFN/JAK1/STAT1 response remained at basal levels in HUVECs regardless of Shoc2 expression levels or its mutation status (**Figure S7B**). Similarly, Shoc2 loss or expression of the Shoc2 S2G variant in primary fibroblasts did not elicit INF response either (**Figure S7C).** These data demonstrate that the aberrant activation of the IFN/JAK1/STAT1 pathway observed following Shoc2 depletion or expression of the S2G variant is unique to HDLECs.

### Activation of RIG-I-like receptor signaling pathway in Shoc2-depleted HDLEC

Additional transcriptome analysis of Shoc2-depleted HDLEC revealed significant enrichment of genes involved in the RIG-I-like receptor (RLR) signaling pathway (KEGG pathway, NES 1.49). Key RLR pathway genes, including *IFIH1* (MDA5), *DDX60*, *DDX58* (RIG-I), *TRIM25*, and *TRIM14*, were significantly upregulated in Shoc2-depleted cells (**Figure 6A**). MDA5 and RIG-I are cytoplasmic pattern recognition receptors that detect viral double-stranded RNA (dsRNA) and trigger an innate immune response. A crucial step in an innate immune response involves the recruitment of MAVS (mitochondrial antiviral signaling protein) located on the outer membrane of mitochondria. The MDA5-MAVS interaction triggers the activation of key transcription factors, including IRF7 and IRF9, which in turn drive the production of IFN and other pro-inflammatory cytokines (**Figure 6B**). (41) qRT-PCR confirmed the increased expression of *MDA5* (*IFIH1*), *RIG-I* (*DDX58*), and other RLR response components (e.g., E3 ligases *TRIM25* and *TRIM14*, and helicase *DDX60*) in Shoc2-depleted HDLEC. Interestingly, *MDA5* and *RIG-I* expression was significantly elevated as early as 3 days post-transfection (**Figure 6C**).

**Figure 6.**
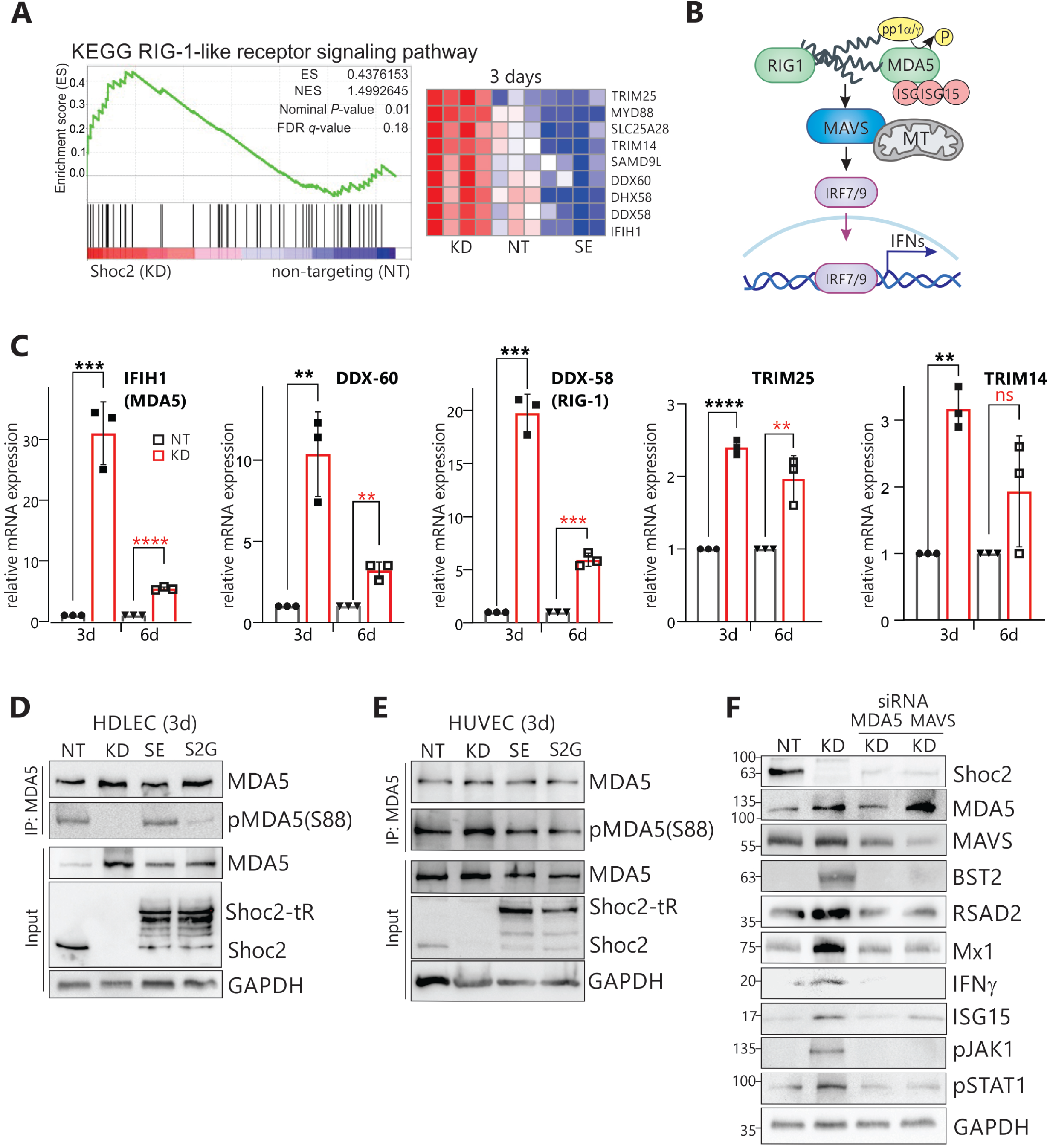
Pattern recognition receptor-mediated response in the HDLEC depleted of Shoc2 or expressing the Shoc2 S2G NSLH variant. **A.** Gene set enrichment analysis (GSEA) is presented as a score plot, with NES indicating the upregulation of selected genes in a RIG-1-like signaling signature in Shoc2-depleted HDLEC (Shoc2 KD, KD) compared with the control (nontargeting, NT). The total height of the curve indicates the extent of enrichment (ES), with the normalized enrichment score (NES), the false discovery rate (FDR), and the p-value. Source data are provided as a source data file. Heatmap: the top elected correlated genes for each biological triplicate are presented in the corresponding heat map. Red and blue indicate increased and decreased expression, respectively, for all heatmaps. **B.** Schematic representation of the RIG-I/MDA5 nucleic acid sensing pathway activated by dsRNA. Activation of MDA5 and its interaction with MAVS leads to phosphorylation of the transcription factors IRF7 and IRF9, ensuring their translocation into the nucleus, where they drive the expression of the IFN genes. **C.** The relative GAPDH-normalized expression of selected genes was measured by RT-qPCR. Data represent the mean and standard error of three independent experiments’ mean (SEM) values. P-values were calculated using a *t*-test (\**p =* 0.01 to 0.05, \*\**p* = 0.001 to 0.01, \*\*\**p =* 0.0001 to 0.001, \*\*\*\**p* < 0.0001, ≥ 0.05-non-significant, ns). Black bars represent control (NT), and red bars represent Shoc2-depleted cells (KD). **D.** MDA5 was precipitated from HDLEC cells depleted of Shoc2 (KD), control (NT), expressing Shoc2-tagRFP (SE), or the Shoc2 S2G variants using anti-MDA5 antibodies. Immunoblots were analyzed with anti-pMDA5 (Ser88), anti-MDA5, and anti-GAPDH antibodies. **E.** MDA5 was precipitated from HUVEC cells depleted of Shoc2 (KD), control (NT), expressing Shoc2-tagRFP (SE), or the Shoc2 S2G variants using anti-MDA5 antibodies. Immunoblots were analyzed using anti-pMDA5 (Ser88), MDA5, and GAPDH antibodies. **F.** Western blot of whole-cell lysates extracted from HDLEC cells expressing lentivirally supplied nontargeting shRNA (NT), Shoc2 shRNA (KD), and transfected with the indicated siRNAs for 3 days. Cell lysates were probed using antibodies to the indicated proteins. The results in each panel are representative of at least three independent experiments.

MDA5 phosphorylation is critical for its activation. (41, 42) Specifically, protein phosphatases PP1cα and PP1cλ-mediated dephosphorylation of Ser88 is essential for activating MDA5 and reversing its signaling-repressed state. (42) Moreover, for MDA5 to translocate to mitochondria and initiate an innate immune response, both dephosphorylation of Ser88 and ISGylation are necessary. (43) We found that 3 days post-transfection increased MDA5 expression (**Figure 6D, input**) correlated with significantly decreased Ser88 phosphorylation on MDA5 precipitated from either Shoc2-depleted or S2G-expressing HDLECs (**Figure 6D, IP**), indicating the MDA5 activation. Yet, Ser88 phosphorylation of MDA5 or its levels remained unchanged in HUVEC cells regardless of Shoc2 levels (**Figure 6E**).

Next, we used MAVS- or MDA5-specific siRNAs to probe whether MDA5 and MAVS drive the IFN response in Shoc2-depleted HDLECs (**Figure 6F**). We found that MDA5 depletion abrogated the IFN response (as assessed by BST2, RSAD2, MX1, and ISG15 expression) and the JAK1/STAT1 pathway activation (as assessed by JAK1 and STAT1 phosphorylation) in Shoc2-depleted HDLEC. siRNA depletion MAVS also markedly reduced the IFN-JAK1/STAT1 response (**Figure 6F**), demonstrating that IFN-JAK1/STAT1 pathway activation in the Shoc2-depleted HDLEC depends on the MDA5-MAVS cytoplasmic pattern recognition response.

### Mitochondrial dysfunction of Shoc2-depleted HDLEC

The outer mitochondrial membrane protein MAVS is involved in maintaining mitochondrial homeostasis and regulating mitochondrial dynamics, including mitochondrial fusion and fission. (44, 45) Hence, we investigated the effects of Shoc2 loss on mitochondrial function. Compared to controls, phosphorylation of mitochondrial fission marker Drp1 (Ser616) was significantly increased in Shoc2-depleted HDLECs and cells expressing the Shoc2 S2G variant (**Figure 7A**). While total DRP1 protein levels remained constant, the levels of other proteins involved in mitochondrial fusion dynamics, including MTFP1, MFF, and OPA1, were altered by the Shoc2 loss. Additional transcriptome analysis uncovered changes in the expression of genes involved in oxidative phosphorylation (OXPHOS), including those encoding ATP synthase, cytochrome c oxidase, and other electron carriers (**Figure S8**).

**Figure 7.**
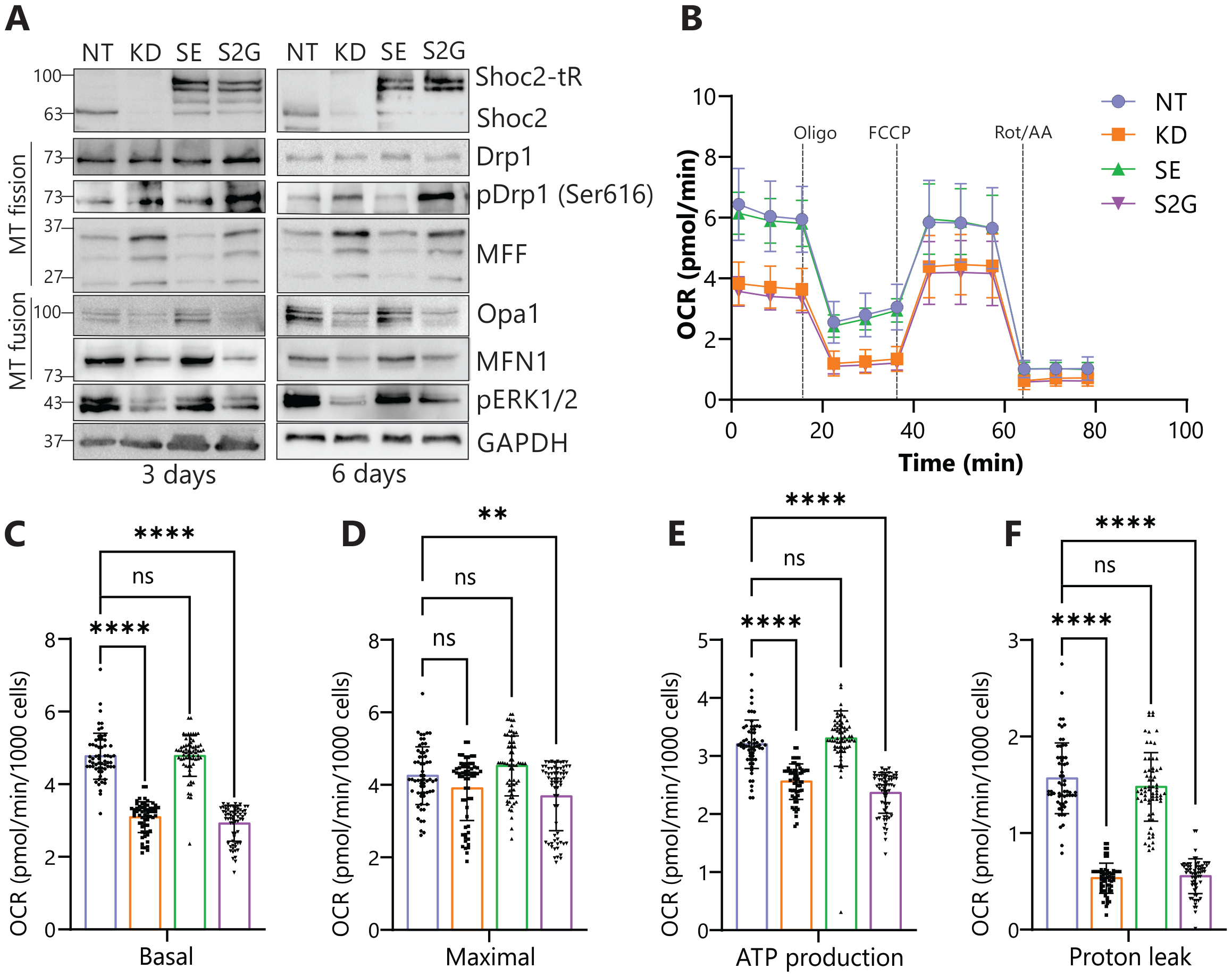
Mitochondrial changes in Shoc2-depleted HDLEC. **A.** Western blot of whole-cell lysates extracted from HDLEC expressing lentivirally supplied nontargeting shRNA (NT), Shoc2 shRNA (KD), Shoc2 shRNA together with Shoc2-tagRFP (SE), or Shoc2 shRNA together with Shoc2(S2G)-tagRFP (S2G) for 3 and 6 days. Cell lysates were probed using antibodies relevant to mitochondrial fission and fusion. GAPDH was used as a loading control. **B.** Relative oxygen consumption rate (OCR), measured by the Seahorse XFe96 analyzer, is normalized to cell number. Data are presented as mean ± SEM of three independent experiments. Statistical significance was determined by one-way ANOVA (brown Forysthe), Dunnett’s multiple comparisons test (\**p =* 0.01 to 0.05, \*\**p* = 0.001 to 0.01, \*\*\**p =* 0.0001 to 0.001, \*\*\*\**p* < 0.0001, ≥ 0.05-non-significant, ns). FCCP, carbonyl cyanide-*p* trifluoromethoxyphenylhydrazone; Rot, rotenone; AA, antimycin A. **C-F.** Basal and maximal OCR analysis, Mitochondrial ATP production, and proton leak of HDLEC depleted of Shoc2 (KD) or expressing either Shoc2-tagRFP (SE) or Shoc2-tagRFP S2G (S2G). Statistical significance was determined by one-way ANOVA (brown Forysthe), Dunnett’s multiple comparisons test (\**p =* 0.01 to 0.05, \*\**p* = 0.001 to 0.01, \*\*\**p =* 0.0001 to 0.001, \*\*\*\**p* < 0.0001, ≥ 0.05-non-significant, ns).

Next, we examined whether mitochondrial respiration was affected by the Shoc2 loss (**Figure 7B**). Seahorse live cell metabolic flux analysis revealed a significant decrease in basal but not maximal consumption rates (OCR) (**Figure 7C, D**). We also found that mitochondrial ATP production (**Figure 7E**) and proton leak (**Figure 7F**) were significantly decreased in cells depleted of Shoc2 or expressing the Shoc2 S2G variant. Together, these data indicate that mitochondrial function is impaired in cells depleted of Shoc2 or expressing the NSLH variant S2G.

### Regulation of the mTOR pathway by Shoc2

The findings that the upregulated IFN-JAK1/STAT1 response is independent of Shoc2’s role in ERK1/2 pathway activation (**Figure 5**) and that there were impairments in mitochondrial function in Shoc2-depleted HDLECs (**Figure 7**) prompted us to investigate other signaling mechanisms potentially dysregulated in Shoc2-depleted HDLECs. Previous work has shown that Shoc2 negatively regulates mTORC1 signaling by competing with mTOR for binding of Raptor, a key mTORC1 complex component (**Figure 8A**). (46) Thus, we examined the effect of Shoc2 loss on mTOR pathway activation in HDLECs (**Figure 8B**). In Shoc2-depleted or S2G variant-expressing HDLECs, increased mTOR phosphorylation correlated with increased phosphorylation of mTOR downstream substrates (i.e., eukaryotic translation initiation factor 4E-binding protein 1 (4E-BP1) and ribosomal protein S6 kinase beta-1 (S6K1)), indicating enhanced mTORC1 activity. Using immunoprecipitation, we showed that Raptor-mTOR binding significantly increased in HDLEC depleted of Shoc2 or expressing the Shoc2 S2G variant. Conversely, the expression Shoc2-tagRFP reduced Raptor interaction with mTOR (**Figure 8C**). Neither the Shoc2 deletion nor the expression affected mTOR-Rictor binding (**Figure 8C, F**).

**Figure 8.**
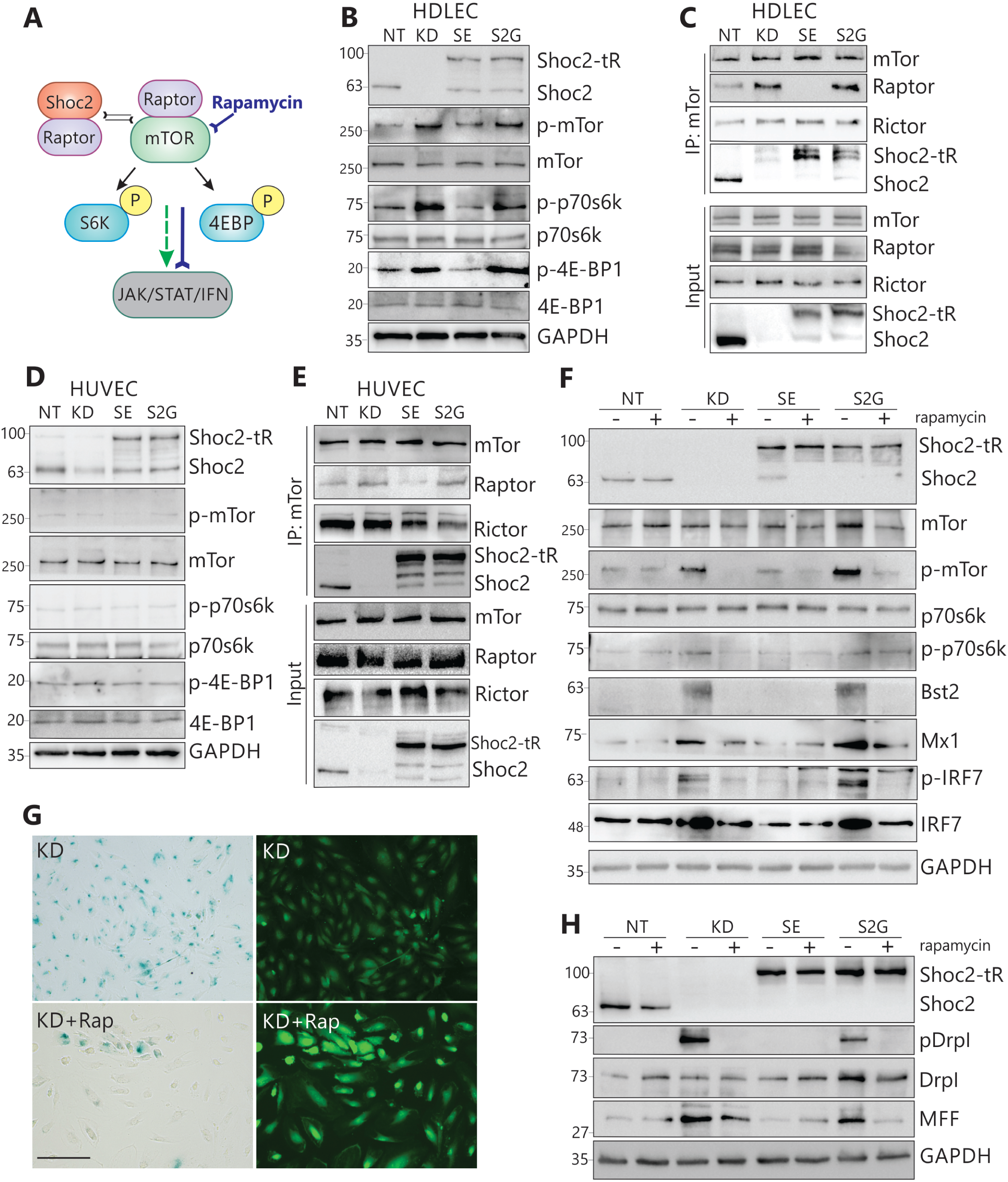
Shoc2 controls mTOR pathway activation in HDLEC. **A.** Schematic representation of the Shoc2-mTOR axis in which Raptor binding to mTOR stimulates mTORC1 complex activity indicated by S6K and 4EBP phosphorylation. mTORC1 activation is followed by activation of the IFN/Jak/STAT response. **B.** Western blot of whole cell lysates extracted from HDLEC expressing lentivirally supplied nontargeting shRNA (NT), Shoc2 shRNA (KD), Shoc2 shRNA together with Shoc2-tagRFP (SE), or Shoc2 shRNA together with Shoc2(S2G)-tagRFP (S2G) for 3 days. Cell lysates were probed using antibodies to proteins relevant to mTORC1 signaling. GAPDH was used as a loading control. **C.** Cell lysates were prepared from HDLEC cells expressing lentivirally supplied nontargeting shRNA (NT), Shoc2 shRNA (KD), Shoc2 shRNA together with Shoc2-tagRFP (SE), or Shoc2 shRNA together with Shoc2(S2G)-tagRFP (S2G) for 3 days. mTOR was immunoprecipitated with an anti-mTOR antibody. Immunoprecipitated proteins were subjected to SDS-PAGE and western blot analysis. Blots were probed with antibodies against mTOR, Raptor, and Rictor. **D.** Western blot of whole cell lysates extracted from HUVEC expressing lentivirally supplied nontargeting shRNA (NT), Shoc2 shRNA (KD), Shoc2 shRNA together with Shoc2-tagRFP (SE), or Shoc2 shRNA together with Shoc2(S2G)-tagRFP (S2G) for 3 days. Cell lysates were probed using antibodies to proteins relevant to mTORC1 signaling. GAPDH was used as a loading control. **E.** Cell lysates were prepared from HUVEC cells expressing lentivirally supplied nontargeting shRNA (NT), Shoc2 shRNA (KD), Shoc2 shRNA together with Shoc2-tagRFP (SE), or Shoc2 shRNA together with Shoc2(S2G)-tagRFP (S2G) for 3 days. mTOR was immunoprecipitated with an anti-mTOR antibody. Immunoprecipitated proteins were subjected to SDS-PAGE and western blot analysis. Blots were probed with antibodies against mTOR, Raptor, and Rictor. **F.** Western blot of whole cell lysates extracted from HDLEC expressing lentivirally supplied nontargeting shRNA (NT), Shoc2 shRNA (KD), Shoc2 shRNA together with Shoc2-tagRFP (SE), or Shoc2 shRNA together with Shoc2(S2G)-tagRFP (S2G) for 3 days and treated with Rapamycin. Cell lysates were probed using the indicated antibodies. GAPDH was used as a loading control. **G.** HDLEC expressing lentivirally-supplied Shoc2 shRNA (KD) were treated with 10nM rapamycin 24 hours after transfection for 6 days and then stained to examine β-Galactosidase activity (left panels). GFP, expressed independently by the lentivirus, served as a reporter to monitor delivery efficiency (right panels). Scale bar – 150μm. **H.** Western blot of whole cell lysates extracted from HDLEC expressing lentivirally supplied nontargeting shRNA (NT), Shoc2 shRNA (KD), Shoc2 shRNA together with Shoc2-tagRFP (SE), or Shoc2 shRNA together with Shoc2(S2G)-tagRFP (S2G) for 3 days and treated with Rapamycin. Cell lysates were probed using the indicated antibodies. GAPDH was used as a loading control.

Interestingly, mTOR pathway hyperactivation was not observed in Shoc2-depleted HUVECs (**Figure 8D**). While mTOR immunoprecipitation showed a slight increase in Raptor-mTOR binding in Shoc2-depleted HUVEC and cells expressing the Shoc2 S2G variant, the densitometric ratio of Raptor to mTOR was significantly lower in Shoc2-depleted HUVEC (1:0.2) than observed in HDLECs (1:1), which may account for the lack of mTOR signaling upregulation in Shoc2-depleted HUVECs (**Figure 8E**).

To confirm the mechanistic link between mTORC1 activation and the IFN response in Shoc2-depleted HDLEC, cells were treated with the mTOR inhibitor Rapamycin. Rapamycin treatment effectively blunted mTOR signaling and blocked the IFN/JAK/STAT response in Shoc2-depleted and S2G-expressing HDLECs (**Figure 8F**). Furthermore, rapamycin treatment also effectively rescued the cell senescence associated with the loss of Shoc2 (**Figure 8G**), as well as changes in activation and expression of proteins involved in mitochondrial fusion dynamics (**Figure 8F**).

These data demonstrate that mTORC1, but not mTORC2, mediates the effect of Shoc2 on IFN signaling in HDLEC and indicate that the Shoc2-Raptor-mTOR equilibrium is critical for maintaining proper levels of mTOR signals in HDLECs. Shoc2 loss or the presence of the S2G mutant disrupts this equilibrium, leading to increased Raptor-mTOR binding. Together, our results suggest that in the absence of Shoc2, hyperactive mTORC1 signaling alters mitochondrial function, which could explain the increased mitochondrial fission, activation of the IFN/JAK/STAT response, and, ultimately, cell senescence (**Figure S9**).

## DISCUSSION

Lymphatic vessel development and homeostasis are complex processes governed by diverse signaling pathways. (1, 5, 47) Our study reveals a novel and essential role for the signaling scaffold Shoc2 in lymphatic regulation, demonstrating its complex influence on cellular signaling and homeostasis. Specifically, Shoc2 loss ablates lymphatic vasculature *in vivo*, while its deficiency in LEC induces (1) activation of mTOR and IFN/JAK/STAT signaling pathways, (2) altered mitochondrial respiration and function, and (3) cellular senescence. Notably, the Shoc2 (S2G) NSLH variant recapitulates the mitochondrial dysfunction, mTOR activation, and IFN/JAK/STAT pathway dysregulation observed in Shoc2-depleted LECs. Thus, this study establishes a novel mechanistic link between the Shoc2 scaffold, mitochondrial function, mTOR signaling, and innate immune response in lymphatic cells. Importantly, these effects are cell-type specific, as Shoc2 depletion in primary fibroblasts and vein endothelial cells did not elicit mTOR pathway activation or immune responses. The differential cellular responses to the loss of this ubiquitously expressed signaling scaffold warrant further investigation to explore whether cell developmental origin, cellular state (proliferating vs. quiescent), or differences in upstream signaling are factors contributing to these cell-specific outcomes.

The embryonic lethality of conventional Shoc2 knockout mice at E8.5, with partial embryo absorption occurring before the onset of lymphangiogenesis, has limited *in vivo* studies. (26) Thus, the Shoc2 mutant zebrafish is currently the sole vertebrate model for examining global Shoc2 loss *in vivo*. Our previous work has demonstrated severe developmental defects in the neural crest of *shoc2* mutant zebrafish. (25, 48) This study highlights the significance of Shoc2 for lymphangiogenesis and demonstrates a Shoc2-autonomous function in lymphatic endothelium (**Figures 1-2**). Shoc2 mutant zebrafish exhibit a striking loss of facial and trunk lymphatic vessels with no apparent effect on other endothelial cells. Given the high homology between zebrafish and human Shoc2, along with ubiquitous expression, these findings are highly relevant to human biology. Our endothelial-specific transgenic Shoc2 “rescue” experiments demonstrate the endothelial-autonomous requirement for lymphatic development and show that the endothelial function of Shoc2 is sufficient for normal lymphatic development. Future studies will investigate whether the tissue microenvironment (e.g., stiffness, immune cell composition) impacts Shoc2 function in different endothelial cells.

### Shoc2 function in lymphatic endothelial cells: mechanisms

Beyond showing Shoc2’s importance in developmental lymphangiogenesis, this study elucidates the molecular mechanisms by which Shoc2 maintains lymphatic endothelial cell homeostasis. Shoc2 function is best understood within the context of the ERK1/2 pathway and its role in mRas-Shoc2-PP1c holoenzyme. (49) Multiple labs, including ours, have demonstrated that Shoc2 regulates the ERK1/2 phosphorylation amplitude, as well as the proliferation and motility of various immortalized or cancer cell types.(50, 51) More recently, Shoc2 has been implicated in regulating mTORC1 signals impacting growth, survival, and tumorigenesis of pancreatic, lung, and liver cancer cells. (20, 52) However, with the exception of mouse neural progenitors, Shoc2 function in primary cells remains largely unexplored. We provide compelling evidence that, in primary lymphatic endothelial cells, Shoc2 regulates VEGF-ERK1/2 activity and balances mTORC1 signals (**Figures 3, 5, and 8**). Consistent with mTORC1’s well-established role in regulating mitochondrial function (i.e., fission, fusion, etc.) and cellular senescence (53–57), we observed that upregulated mTORC1 in the Shoc2-depleted cells leads to mitochondrial dysfunction. Thus, given mTORC1’s pleiotropic roles in cellular processes, such as nutrient/energy sensing, protein synthesis, and metabolism, additional work to fully elucidate the spectrum of Shoc2’s effects on cellular homeostasis, particularly its role in adaptive autophagy and nutrient sensing, will be critical.

In this study, we also provide new evidence that Shoc2 dysfunction impacts mitochondria-regulated innate immune response (**Figure 7**). Mitochondria function as both energy providers and signaling hubs in the innate immune response. To regulate immune responses, mitochondria release signals like mitochondrial DNA (mtDNA), RNA (mtRNA), and reactive oxygen species (ROS). (45, 58) Our findings of increased expression and activation of MAVS and the pattern recognition receptors, RIG1 and MDA5 (**Figure 6**), suggest that hyperactive mTORC1 induces mitochondrial dysfunction of Shoc2-depleted LECs. While mitochondrial dsRNA is the likely trigger of the response in Shoc2-depleted cells, additional studies are needed to confirm this. Yet, it is reasonable to assume that in the absence of Shoc2, mt-dsRNA activates RIG1/MDA5, triggers MAVS and IRF-7/9 signaling, leading to proinflammatory cytokine and interferon production, and ultimately contributes to cellular stress and senescence (**Figure S9**). Importantly, our data indicate that Shoc2’s role in MDA5 activation is independent of the Shoc2-PP1c-MRas holoenzyme, as PP1c was absent from mTOR-Shoc2 immunoprecipitates and MDA5 Ser338 was dephosphorylated in Shoc2-depleted cells. Whether the exceptional sensitivity of lymphatic endothelium to Shoc2 dysfunction is related to the requirement to limit mTORC1 signals to maintain cell quiescence will also be explored in the following investigations.

The IFNγ inhibitory effects on lymphatic vessels are well-documented. (1, 59, 60). IFNγ treatment reduces the expression of the key lymphatic transcription factor PROX-1 and the membrane glycoprotein LYVE-1 and suppresses lymphatic growth *in vitro*. (59–61) Given the observed increase in IFNγ signaling in Shoc2-depleted cells, it is plausible that IFNγ contributes to their senescence. Moreover, IFNγ’s precise influence on lymphangiogenesis may depend on factors such as other cytokines or the specific type of lymphatic endothelial cell involved. Future studies will determine if elevated IFNγ levels in lymphatic progenitor cells underlie the lymphatic vessel defects observed in Shoc2 knockout larvae. Considering the emerging role of lymphatic cells in shaping local immune responses (62), it is essential to investigate whether lymphatic cells in NSLH patients modulate adaptive immunity or antigen presentation. The physiological consequences of the Shoc2 S2G-associated IFN response on lymphatic vessel development and the potential contribution of chronic IFN activation to lymphatic dysfunction also warrant thorough investigation. Such studies will be in line with the recent findings demonstrating Shoc2’s involvement in the immune signaling of invertebrate antibacterial and eukaryotic antiviral responses. (63, 64)

### Implications for NSLH pathology

One of the most interesting observations of this study is that, in lymphatic cells, the Shoc2 NSLH variant S2G fails to compete effectively with mTOR for Raptor binding. Similarly to the Shoc2 loss, this Shoc2 variant disrupts mitochondrial function and triggers the IFN response. The Shoc2 S2G variant localizes to the PM due to aberrantly acquired *N*-myristoylation. (10) Thus, it is tempting to speculate that these Shoc2-Raptor-mTOR complexes are spatially defined. However, the molecular details of why this variant, despite being able to bind mTOR, still causes increased mTOR activity remain to be elucidated. Notably, mitochondrial dysfunction, decreased adipogenesis, fat malabsorption, increased energy expenditure, and enhanced insulin signaling are found in other RASopathies. (65–67) For instance, PTPN11 mutations causing Noonan syndrome reduce mitochondrial membrane potential and ATP content while increasing ROS levels. (68–70) Similarly, B-RAF and NF1 mutations affect mitochondrial respiration. While the specific mechanisms and downstream effectors contributing to mitochondrial dysfunction in different RASopathies are still unclear, our findings underscore the need for further research to define the mechanistic link between mitochondrial dysfunction and mutations in specific RASopathy genes.

Intriguingly, RASopathy patients, including those diagnosed with NSLH, exhibit an increased risk of autoimmune disorders. (71–73) Individuals with the Shoc2 S2G variant exhibit autoimmune manifestations such as systemic lupus erythematosus, autoimmune cytopenia, and autoimmune thyroiditis. (71, 74–77) Despite this association, the triggers for clinically overt disease remain poorly understood. Therefore, our discovery of a link between mitochondrial dysfunction, IFN response, and cellular homeostasis in Shoc2-deficient lymphatic cells potentially provides a novel mechanism underlying the autoimmune presentation in NSLH patients.

In conclusion, we have established that the progressive IFN response observed in both Shoc2-deficient cells and cells expressing the NSLH S2G variant depends on upstream mTOR signaling and can be abrogated by treatment with the mTOR inhibitor rapamycin. Our findings underscore the critical role of Shoc2 as a molecular gatekeeper that balances ERK1/2 and mTOR signals, highlighting its cell-type-specific functions. We also provide novel insights into the roles of ERK1/2 and mTOR during developmental lymphangiogenesis, opening new avenues for understanding how these pathways contribute to lymphatic development and related disorders. These results raise critical questions for future research and emphasize the need for a deeper understanding of Shoc2’s multifaceted role in healthy and diseased organisms. Therapies combining ERK1/2-targeting therapies with mTOR inhibition may be relevant for treating RASopathies. Given the intricate interplay between Shoc2, IFN response, and mTORC1, targeting these pathways holds significant therapeutic potential for diseases associated with lymphatic dysfunction or aberrant immune activation. Specifically, JAK or mTOR inhibitors may offer therapeutic benefits for patients with Shoc2 mutations, potentially mitigating the observed lymphatic and immune-related pathologies.

## MATERIALS AND METHODS

### Antibodies and other reagents

**Table S1** lists the specific primary and secondary antibodies used for protein detection. Ruxolitinib and PD98059 were purchased from LC Labs, and hexadimethrine bromide (polybrene) was obtained from Sigma-Aldrich.

### Constructs

tagRFP-tagged Shoc2 (Shoc2-tRFP), the pLVTHM constructs expressing Shoc2 short hairpin RNA (shRNA), and nontargeting shRNA were described previously. (78) The pLVTHM constructs expressing Shoc2 and nontargeting shRNA also express GFP to visualize transfected cells. (78)

### Cell culture and lentivirus transduction

Human dermal lymphatic endothelial cells (HDLEC) (PromoCell) and normal human umbilical vein endothelial cells (HUVEC) (Lifeline Cell Technology) were grown in VascuLife basal media (Lifeline Cell Technology) containing 5 ng/mL rh FGF basic, 50 µg/mL ascorbic acid, 1 µg/mL hydrocortisone hemisuccinate, 2% FBS, 10 mM L-glutamine, 15 ng/mL rh IGF-1, 5 ng/mL rh EGF, 5 ng/mL rh VEGF, 0.75 U/mL heparin sulfate, and supplemented with 30 µg/mL gentamicin and 15 ng/mL amphotericin B.

For lentivirus transduction, cells were plated in 12-well dishes at 50–60% confluence. The ratio of viral particles to target cells was determined for each viral prep. The lentiviral stock was diluted in a fresh culture medium containing polybrene. Shoc2–tagRFP (SE or S2G) and shShoc2/GFP (KD), or non-targeting-GFP (NT) fluorescence, were detected 36–48 h after lentivirus transduction. The efficiency of shRNA knockdown and cDNA expression was validated using Shoc2-specific antibodies 3 days after lentivirus transduction.

### Real-time quantitative PCR (qPCR)

Total RNA was isolated from cells using a PureZOL/Aurum total RNA isolation kit (Bio-Rad) according to the manufacturer’s instructions. Aliquots containing equal amounts of RNA were subjected to reverse transcription-PCR (RT-PCR) analysis. Quantitative RT-PCR was performed using SoAdvanced SYBR Green Supermix and a Bio-Rad CFX detection system (Bio-Rad). Sequence-specific primer sets are listed in **Table S2**. HPRT1 mRNA was used as a reference gene. The relative amounts of RNAs were calculated using the comparative threshold cycle method. The values for the samples were normalized against those for the reference gene, and the results are presented as the fold change in the amount of mRNA recovered from cells transfected with nontargeting siRNA. The data represent the mean standard deviations (SDs) from two independent experiments.

### siRNA transfections

To silence protein expression by RNA interference, cells were seeded in 12-well plates (at 50 to 60% confluence with 1 ml of VascuLife media per well) at least 20 h before transfection. Small interfering (siRNA) transfections were performed according to the manufacturer’s recommendations, using Dharmafect reagent 2 (Dharmacon). The siRNA sequences used to target MDA5 and MAVS are listed in **Table S4.** The efficiency of the siRNA knockdown was validated by western blotting.

### Immunoprecipitation

Cells were placed on ice and washed with Ca^2+^, Mg^2+^-free phosphate buffered saline (PBS), and the proteins were solubilized in 50 mM Tris (pH 7.5) containing 150 mM NaCl, 1% Triton X-100, 1 mM Na_3_VO_4_, 10 mM NaF, 0.5 mM phenylmethylsulfonyl fluoride (PMSF, Sigma, St. Louis, MO, USA), 10 μg/ml of leupeptin, and 10 μg/ml of aprotinin (Roche, Basel, Switzerland) for 15 min at 4°C. Lysates were centrifuged at 14,000 *rpm* for 15 min to remove insoluble material. Lysates were incubated with appropriate antibodies for 2 hrs, and the immunocomplexes were precipitated using protein A- or G-Sepharose (GE Healthcare Life Sciences, Chicago, IL, USA). Immunoprecipitates and aliquots of cell lysates were denatured in a sample buffer at 95°C, resolved by electrophoresis by SDS-PAGE, transferred to a nitrocellulose membrane, and probed by western blotting with various antibodies, followed by chemiluminescence detection.

mTOR immunoprecipitation was done following the protocol of Kim et al. 2002. Briefly, cells were rinsed once with PBS, lysed in 600 µl of ice-cold lysis buffer (40 mM HEPES [pH 7.5], 120 mM NaCl, 1 mM EDTA, 50 mM NaF, 1.5 mM Na3VO4, 0.5 mM phenylmethylsulfonyl fluoride, 10 μg/ml of leupeptin, and 10 μg/ml of aprotinin and 0.3% CHAPS) and incubated for 30 min on ice. After clearing, 40 µl of a 50% slurry of protein G-Sepharose and 4 µg of the anti-mTOR antibody were added to the supernatant and rotated for 3 hrs at 4 °C.

### Western blot analysis

Cells were placed on ice and washed with Ca^2+^, Mg^2+^-free phosphate buffered saline (PBS), and the proteins were solubilized in 50 mM Tris (pH 7.5) containing 150 mM NaCl, 1% Triton X-100, 1 mM Na_3_VO_4_, 10 mM NaF, 0.5 mM phenylmethylsulfonyl fluoride (PMSF, Sigma, St. Louis, MO, USA), 10 μg/ml of leupeptin, and 10 μg/ml of aprotinin (Roche, Basel, Switzerland) for 15 min at 4°C. Lysates were centrifuged at 14,000 *rpm* for 15 min to remove insoluble material. Aliquots of cell lysates were denatured in a loading buffer at 95°C, resolved by electrophoresis, and probed by western blotting with various antibodies, followed by chemiluminescence detection. Proteins transferred from SDS-polyacrylamide gels to nitrocellulose membranes were visualized using a ChemiDoc analysis system (Bio-Rad, Hercules, CA, USA). Several exposures were analyzed to determine the linear range of the chemiluminescence signals. Quantification was performed using the densitometry analysis mode of Image Lab software (Bio-Rad, Hercules, CA, USA). To visualize proteins with close molecular weight, in some instances, several identical SDS-PAGE gels were used to resolve protein lysates.

### Seahorse extracellular flux analysis

The Seahorse XF96 Extracellular Flux Analyzer (Agilent) was used to measure cellular respiration activity in HDLEC. Cells were seeded at a density of 2×10^4^ cells per well in an XF96 plate ∼12 h before the measurement. The following day, cells were washed twice with assay medium (unbuffered DMEM supplemented with 25 mM glucose, 2 mM L-glutamine, and 1 mM pyruvate) and incubated at 37°C in a non-CO2 incubator for 1 hour. OCR was measured at baseline and following sequential injections of oligomycin (1 µM), carbonyl cyanide-4-(trifluoromethoxy)phenylhydrazone (FCCP) (1.5 µM), and rotenone/antimycin A (0.5 µM). Data were normalized to total protein content determined by a BCA assay.

The mitochondrial stress tests were performed per the manufacturer’s protocol. The relative levels of non-mitochondrial, basal, maximal, and ATP production-related respiration were calculated using the Mito stress tests using the Seahorse Wave software (Agilent) for XF analyzers.

### Cell senescence assay

HDLEC were stained for senescence-associated β-galactosidase activity using the β-Galactosidase Staining Kit (Cell Signaling) according to the manufacturer’s recommendation. Briefly, equal numbers of cells 2×10^4^ were washed once with 1X PBS and then fixed. Cells were incubated with the β-Galactosidase Staining Solution ON at 37°C for at least overnight. Cells were overlaid with 70% glycerol for long-term storage and stored at 4°C. To quantify cells stained for senescence-associated β-galactosidase activity, we acquired GFP fluorescence to visualize the transfected cells. The percentage of senescent cells was divided by the total number of cells counted using immunofluorescence (*n* = 3 dishes per genotype).

### Zebrafish strains and maintenance

This study was conducted in AAALAC-accredited facilities under active research projects approved by the University of Kentucky Animal Care and Use Committee (#2019-3208), the University of Illinois at Chicago Animal Care and Use Committee (#23-112), and the *Eunice Kennedy Shriver* National Institute of Child Health and Human Development (Animal Study Proposal #21-015, 24-015). All zebrafish strains were bred, raised, and maintained according to established animal care protocols for zebrafish husbandry. Embryos were staged as previously described. (79) The following transgenic and mutant zebrafish lines were used in this study: *Tg(mrc1a:eGFP*)*^y251^*(^28^)*, Tg(kdrl:mcherry)^y205^*(80), and *Tg(fli1:eGFP)^y1^*.(81) The *Tg(mrc1a:KalTA4)* line was generated from this study. The *Tg(UAS:eGFP)* line was generously provided by the Kawakami lab. The *shoc2^Δ22^* has been previously described in (25), and the *shoc2^SA/SA^* line was obtained from the Sanger Institute Zebrafish Mutation Project.

### Generation of transgenic zebrafish lines

The *mrc1a:KalTA4* transgene construct was assembled using an LR recombination reaction, using the destination vector *pDestTol2pA/pA2* (82) and three entry clones: *p5E-mrc1a* (28), *pME-KalTA4* (PCR-amplified from plasmid #10080, Addgene), and *p3E-polyA* from the Tol2kit. (82) The pME-KalTA4 construct was amplified using primers containing attB1/attB2 sites, and then cloned into the pDONR211 vector using BP Clonase (Cat# 11789-020, Thermo Fisher Scientific). The transgenic line *Tg(mrc1a:KalTA4)* was generated by microinjecting the *mrc1a:KalTA4* transgene construct (55 pg) along with *Tol2* transposase (50 pg) into one-cell stage zebrafish embryos from the *Tg(UAS:eGFP)* line. Embryos were subsequently screened for GFP expression in the lymphatic vasculature, leading to the establishment of the germline-transmitting line *Tg(mrc1a:KalTA4;UAS:eGFP) ^y721^*.

### Imaging methods

Embryos were anesthetized using 1x tricaine, mounted in 0.8-1.5% low-melting-point agarose dissolved in egg water, and mounted on a depression slide. Confocal images were acquired using a Nikon Ti2 inverted microscope with a Yokogawa CSU-W1 spinning disk or a Zeiss LSM880 microscope.

Brightfield images were obtained using a Leica M205 microscope. The images were analyzed using ImageJ, Adobe Photoshop (Adobe), and NIS-Elements (Nikon) software.

### Morpholino injection

All MOs were obtained from Gene Tools, LLC (Philomath, OR) and injected into 1-2 cell stage zebrafish embryos. The following MOs were used in this study: standard control MO: 5’-CCTCTTACCTCAGTTACAATTTATA-3’; *shoc2* MO1: 5’-TACTGCTCATGGCGAAAGCCCCGCA-3’. Embryos were injected with 5.2 ng of Shoc2 MOs.

### Shoc2 expression construct

The 6x-UAS:mCherry-2A-shoc2 construct was generated through a three-part LR recombination reaction using Gateway Technology. This process involved the creation of three entry clones and a destination vector. The 5’ entry clone (p5E 6xUAS) was constructed via traditional cloning methods. A linearized p5E-MCS vector, digested with XhoI and SpeI, served as the backbone, into which a 6xUAS fragment was inserted. This fragment was PCR-amplified from the 6x-UAS:EGFP plasmid, kindly provided by Marnie Halpern. The middle entry clone (pME mCherry-2A-shoc2) was generated using SLiCE (<u>S</u>eamless <u>Li</u>gation <u>C</u>loning <u>E</u>xtract, where 3 fragments were assembled by short stretches of homologous DNA recombination with the PPY SLiCE strain. The first fragment was a linearized pME-MCS vector, digested with EcoRI and SpeI. The second fragment was a PCR-amplified mCherry-2A fragment from a plasmid encoding mCherry-2A, and the third fragment was a full coding sequence of zebrafish shoc2 amplified using PCR. The 3’ entry clone, p3E polyA, containing the polyA sequence, was obtained from the Tol2kit. The final 6xUAS mCherry-2A-shoc2 construct was assembled by performing an LR recombination reaction using the destination vector, pDestTol2pA/pA2 (82), resulting in the insertion of the three entry clones into the destination vector.

### Shoc2 rescue in zebrafish

Shoc2 rescue experiments were conducted in shoc2 morphants with the *Tg(mrc1a:kalTA4; UAS:eGFP)* transgenic background for endothelial rescue. The shoc2 MO (5’-TACTGCTCATGGCGAAAGCCCCGCA-3’; Gene Tools, LLC) was used at a concentration of 4.6 ng per embryo. For Shoc2 expression rescue, embryos were injected with either 6x-UAS:mCherry-2A-shoc2 or 6x-UAS:mCherry (control group) at a concentration of 40-45 ng per embryo, with or without shoc2 MO. DNA constructs were injected at the one-cell stage, followed by MO injection into the yolk at the one-to-four-cell stage. Image acquisition was performed using a Nikon Ti2 inverted microscope with Yokogawa CSUW1 spinning disk confocal, with the 10× Air 0.45 NA objective.

### Analysis of RNA-seq data

mRNAs were isolated from HDLECs expressing Shoc2 shRNA. For library preparation, mRNA was first extracted from total RNA using oligo (dT) magnetic beads and sheared into short fragments of about 200 bases. The cDNA library was sequenced using the Illumina NextSeq 500 sequencer. Quality control (QC) of the raw sequence data was performed using FastQC (version 0.11.7). The concatenated sequences were directly aligned to the Homo Sapience GRCh38 reference genome assembly (GRCh38) using STAR (version 2.6), generating alignment files in bam format. Sequencing reads were trimmed and filtered using Trimmomatic (V0.39) to remove adapters and low-quality bases. Trimmed reads were mapped to the human reference genome assembly GRCh38 transcripts annotation using RSEM. RSEM results normalization and differential expression analysis were performed using the R package EdgeR. Significantly up-/downregulated genes between groups were determined as fold change >= 2 and q-value <0.05. Transcript per million were used for generating heatmaps, with represented values ranging from +1.5 (red, high expression) to −1.5 (blue, low expression). The RNA-seq data is publicly available as GEO series GSE288882. The data is MIAME compliant (https://www.ncbi.nlm.nih.gov/geo/query/acc.cgi?acc=GSE288882).

The gene set enrichment analysis was performed using GSEA software and the gene sets in the Molecular Signature Database (MSigDB). As recommended for GSEA from RNA sequencing experiments, genes with no counts in any sample were removed, along with low-expressing genes (mean or geometric mean <10 reads across all samples). As recommended in the GSEA user guide (https://www.gsea-msigdb.org/gsea/doc/GSEAUserGuideFrame.html), a false discovery rate (FDR) cut of 25% was used for Hallmark, KEGG, and Biological pathway gene set analysis. These data were the basis for the Venn diagram, heatmaps, and KEGG pathway analysis.

### Statistical analyses

Results are expressed as means ± S.D. The statistical significance of the differences between groups was determined using a *t*-test or one-way and two-way ANOVA (followed by Dunnett’s test). *p* 0.01 to 0.05 was considered statistically significant. (\**p =* 0.01 to 0.05, \*\**p* = 0.001 to 0.01, \*\*\**p =* 0.0001 to 0.001, \*\*\*\**p* < 0.0001, ≥ 0.05-non-significant, ns). GraphPad Prism 8.0 (GraphPad Prism Software Inc., Chicago, IL, USA) was used to perform all statistical analyses.

## Data availability

This published article and its supporting information files include all data generated or analyzed during this study.

## Supporting information

This article contains supporting information.

## Acknowledgments

We thank Dr. Tianyan Gao for critically reading the manuscript and for valuable reagents, and Dr. Saurabh Chattopadhyay and Dr. Michaela Guck for valuable reagents. We also thank Ms. Lina Abdelmoti for the excellent care of the zebrafish colony.

