## Supplemental data for "Signaling scaffold Shoc2 regulates lymphangiogenesis by suppressing mTORC1-mediated IFN responses"

#### SUPPORTING DATA METHODS AND MATERIALS

##### Materials Availability

All antibodies, chemicals, and zebrafish lines used in this study are available commercially or upon request.

##### Data and code availability

Accession IDs provided by GEO: GSE288882

##### Zebrafish strains and maintenance

All zebrafish (*Danio rerio*) strains were bred, raised, and maintained following established animal care protocols for zebrafish husbandry. Embryos were staged as previously described (1). All animal procedures were carried out per guidelines established by the University of Kentucky Institutional Animal Care and Use Committee. Briefly, zebrafish embryos were raised at 28.5°C, kept in a 14/10h light/dark cycle, and staged according to Kimmel *et al*., 1995. (1) When necessary, 1-Phenyl-2-thiourea (0.002%) was added to the embryo media to prevent pigment development.

*Shoc2*$^{\Delta 22}$ and *Shoc2*$^{SA/SA}$ zebrafish were maintained as heterozygotes and incrossed to generate homozygous mutant embryos. The *Shoc2*$^{SA/SA}$ heterozygous mutant line was crossed with available reporter lines Tg*(mrc1a:eGFP,* Tg*(kdrl1:mCherry)*. The mutant *shoc2* zebrafish line Shoc2 E2Δ22+/- (ZDB-GENE-050208-523) was reported previously (2).

##### Genotyping

Genomic DNA was extracted from individual embryos or adult tail clips. Briefly, 20 µl of the ThermoPol Buffer (New England Biolabs, # B9004S) was added to the samples and boiled for 5 min. Samples were digested with 50µg (5µL) Proteinase K (Millipore Sigma, #p22308) for 12 hours at 55 °C. Proteinase K was then inactivated by boiling for 10 minutes. PCR was carried out in a 25 µl reaction solution containing 1 µl of 10 mM dNTP, 1 µl of 10 mM forward and reverse primer, 2.5 µl of 1x ThermoPol buffer, and 0.5 units of Taq Polymerase (New England Biolabs, #M0267). The primers used to detect heterozygous mutant alleles are listed in **Supplemental Table 3**.

##### Phenotype analysis

##### *Skeletal stain*

For Alcian blue staining, zebrafish larvae were fixed in 4% paraformaldehyde for 2 h at room temperature and stained according to Kimmel et al., 1998. (3) Calcified structures were examined by acid-free Alizarin Red S staining. Larvae were fixed in 4% PFA for 2 h and stained in a 0.05% Alizarin Red S solution for 30 min in the dark on low agitation. Larvae were then rinsed in a 50% glycerol, 0.1% KOH solution to remove excessive staining and kept at 4◦C in the same solution for imaging.

##### *Iridophores*

Incident light images of four-day-old embryos' iridophores were captured with a Leica M165FC microscope. All iridophores in a 1350-um-long region in the tail (spanning approximately 11 somites) were quantified.

##### *Imaging methods and analysis*

Images of whole-body Alcian blue and Alizarin Red stained larvae were acquired using a Leica DFC450 digital camera. Alcian blue ceratohyal images were acquired with a Zeiss Imager AzioCam MRm.

#### Molecular analysis

##### *Real-time quantitative polymerase chain reaction (RT-qPCR)*

Total RNA was isolated from cells or a pool of 25 embryos using PureZOL RNA Isolation Reagent (Bio-Rad, #732-6890) and Aurum Total RNA Isolation Kit (Bio-Rad, #732-6820). Aliquots containing equal amounts of RNA were subjected to RT-PCR analysis (Bio-Rad, iScript™ Reverse Transcription Supermix for RT-qPCR, # 1708840). qPCR was performed using Bio-Rad iTaq™ Universal SYBR® Green Supermix (#1725120) and a Bio-Rad CFX detection system (Bio-Rad, CA). Relative amounts of RNAs were calculated using the comparative $C_T$ method. Sequence-specific primer sets are presented in **Supplemental Table 2**.

##### *Western blot analysis*

Proteins were extracted from dechorionated and de-yolked embryos/larvae and resolved by SDS-PAGE. In short, water was removed from approximately 25 embryos in a microcentrifuge tube. One solid glass bead and 50 μL RIPA buffer containing protease inhibitors were added to the embryos. Gentle manual agitation physically lysed the embryos. Samples were centrifuged at 4 °C for 10 minutes at 14,000 RPM. Total protein lysates were removed from the pellet and beads. 25μg of total lysate per sample was resolved on a 10% acrylamide gel. Western blot analysis was performed as described previously. (4) Quantification was performed using the densitometry analysis mode of Image Lab software (Bio-Rad, CA). The antibodies used in these experiments are listed in

#### SUPPORTING DATA FIGURE LEGENDS

**Figure S1. Molecular analysis of zebrafish larvae carrying *shoc2*$^{SA}$ allele.**

**A.** To detect *shoc2* WT and Shoc2$^{SA}$ mutant alleles in individual larvae, genomic DNA was used to amplify a DNA fragment that was subsequently digested by HphI digestion. The *shoc2*$^{SA}$ allele c.1546 G>A substitution alters the HphI recognition sequence 5'…GGTGA(N)$_8$…3', eliminating the HphI restriction site. Agarose gel electrophoresis shows PCR amplicons of WT, heterozygous, and homozygous *shoc2*$^{SA}$ allele before and after HphI digest. Primers used in these experiments are listed in **Table S2**

**B.** cDNA fragment amplified using mRNA isolated from *shoc2*$^{SA/SA}$ mutants was longer than a similar fragment amplified from WT larvae. The *shoc2*$^{SA/SA}$ allele incorporates 31 intronic nucleotides into the mRNA transcript, resulting in a longer amplicon (289 nucleotides) than WT's fragment (258 nucleotides). Agarose gel electrophoresis shows amplicons of WT and *shoc2*$^{SA/SA}$ alleles.

**C.** Schematic representation of the Shoc2 loci, gene, and alternative splicing site. The results of sequence analysis of cDNA amplified using *shoc2*$^{SA/SA}$ mRNA. The sequencing result confirms the reported mutation. The asterisk indicates the G>A point mutation.

**D.** The relative *ath5*-normalized expression of Shoc2 was measured by RT-qPCR using mRNA extracted from 6 dpf WT and *shoc2*$^{SA/SA}$ larvae. The data are presented as the fold change of the mRNA levels in WT larvae versus those in *shoc2*$^{SA/SA}$ larvae larvae. The results represent an average of three biological replicas. Error bars indicate means with SEM. ****P<0.0001 at 95% confidence interval (two-tailed t-test).

**E.** Western blot analysis of Shoc2 WT and *shoc2*$^{SA/SA}$ larvae. Larvae were harvested at 6 dpf. Protein expression was analyzed using specific antibodies. β-actin was used as a loading control. The results represent an average of three biological replicas.

**F.** *shoc2*$^{SA/SA}$ larvae develop edema. Edemic larvae were collected and genotyped at 4 dpf (light grey), 5 dpf (dark grey), and 6 dpf (black) (as shown on the right axis). All remaining embryos were collected and genotyped at 6 dpf. N represents the number of genotyped larvae.

**G.** PCR analysis of genomic DNA of *shoc2*$^{+/SA}$ inbred larvae at 11 dpf. Progeny from three independent experiments using heterozygous *shoc2*$^{+/SA}$ larvae were genotyped. All surviving larvae were *shoc2* WT or heterozygous carriers of the *shoc2*$^{SA}$ mutant allele.

**Figure S2. Developmental analysis of the *shoc2*$^{SA}$ allele.**

**A.** Lateral and ventral view of 6 dpf WT or *shoc2*$^{SA/\Delta22}$ larvae stained with Alcian blue. Mutant larvae show significant changes in head cartilage. The Meckel's Cartilage was dissected and mounted before imaging to highlight the hypoplastic closure in *shoc2*$^{SA/\Delta22}$ larvae.

**B.** Lateral view of a 6 dpf WT or *shoc2*$^{SA/\Delta22}$ larvae stained with Alizarin Red S. *shoc2*$^{SA/\Delta22}$ larvae show significant differences in cranial bone formation.

**C.** Iridophores from 6 dpf WT *shoc2*$^{SA/SA}$ larvae were detected using iridescent light.

**D.** Iridophores of WT and *shoc2*$^{SA/\Delta22}$ larvae were quantified from three biological replicas. ***P<0.001 at 95% confidence interval (two-tailed t-test).

**E.** Dorsal (head) and lateral (trunk) views of 6 dpf WT and *shoc2*$^{SA/\Delta22}$ larvae. Unlike WT larvae, *shoc2*$^{SA/\Delta22}$ larvae present closed gaps in the pigmentation patterning of both the head and lateral stripe (see inset).

**Figure S3. Primary sprouts are normal in shoc2 mutant zebrafish.**

**A.** Confocal image of trunk vessels of a wild-type sibling *Tg(fli1a:eGFP)*$^{y1}$ zebrafish larvae at 24 hpf. Scale bar: 100 µm.

**B.** Confocal image of trunk vessels of a wild-type sibling *Tg(fli1a:eGFP)*$^{y1}$ zebrafish larvae at 27 hpf. Scale bar: 100 µm.

**C.** Confocal image of trunk vessels of a wild-type sibling *Tg(fli1a:eGFP)*$^{y1}$ zebrafish larvae at 30 hpf. Scale bar: 100 µm.

**D.** Confocal image of trunk vessels of a *shoc2*$^{SA/SA}$ mutant *Tg(fli1a:eGFP)*$^{y1}$ zebrafish larvae at 24 hpf. Scale bar: 100 µm.

**E.** Confocal image of trunk vessels of a *shoc2*$^{SA/SA}$ mutant *Tg(fli1a:eGFP)*$^{y1}$ zebrafish larvae at 27 hpf. Scale bar: 100 µm.

**F.** Confocal image of trunk vessels of a *shoc2*$^{SA/SA}$ mutant *Tg(fli1a:eGFP)*$^{y1}$ zebrafish larvae at 30 hpf. Scale bar: 100 µm.

**Figure S4. Transcriptome profiling of data sets in Figure 4.**

**A.** Western blot analysis of human dermal lymphatic endothelial cells (HDLEC) expressing non-targeting (NT) shRNA, Shoc2 shRNA(KD), or expressing Shoc2-tagRFP (SE). Cell lysates were analyzed using anti-pERK1/2, -ERK1/2, and -Shoc2 antibodies.
**B.** Venn diagram showing the number of differentially expressed genes in HDLECs depleted (KD) of Shoc2 or expressing Shoc2-tRFP (SE).
**C.** GSEA Hallmark analysis of selected differentially regulated pathways in Shoc2-depleted HDLECs.

**Figure S5. Transcriptome analysis of data sets in Figure 4.**
**A-B.** Gene set enrichment analysis (GSEA) of GOBP data sets presented as a score plot with NES showing upregulation of selected genes of the 'Defence response to virus' (A) and the 'Negative regulation of viral process' (B) signature in Shoc2-depleted HDLEC (Shoc2 KD, KD) compared with control (non-targeting, NT) or HDLEC depleted of endogenous Shoc2 and expressing Shoc2-tRFP (SE). The total height of the curve indicates the extent of enrichment (ES), with the normalized enrichment score (NES), the false discovery rate (FDR), and the p-value. Source data are provided as a source data file.
**C.** Heatmap showing differentially expressed IFN alpha genes in HDLEC expressing non-targeting shRNA, Shoc2 shRNA, or Shoc2-tRFP (p-value ≤ 0.05, [$Log^2$fold change] ≥0.0).
**D.** Heatmap showing differentially expressed IFN gamma genes in HDLEC expressing non-targeting shRNA, Shoc2 shRNA, or Shoc2-tRFP (p-value ≤ 0.05, [$Log^2$fold change] ≥0.0).
Red and blue indicate increased and decreased expression for all heatmaps, respectively.

**Figure S6. Analysis of Shoc2-depleted cells.**
**A.** The relative GAPDH-normalized expression of selected genes was measured by RT-qPCR. Data represent the mean and standard error of three independent experiments' mean (SEM) values. P-values were calculated using t-test (**p* = 0.01 to 0.05, ***p* from 0.001 to 0.01, ****p from* 0.0001 to 0.001, *****p* < 0.0001, ≥ 0.05- non-significant, ns)

**Figure S7. Analysis of Shoc2-depleted primary human.**
**Figure 6**. **Cell-specific interferon response in Shoc2-depleted or expressing the Shoc2 S2G NSLH variant HDLEC.**

**A.** Western blot of whole cell lysates extracted from human dermal lymphatic endothelial cells (HDLEC) expressing lentivirally supplied nontargeting shRNA (NT), Shoc2 shRNA (KD), Shoc2 shRNA together with Shoc2-tagRFP (SE), or Shoc2 shRNA together with Shoc2(S2G)-tagRFP for days. Cell lysates were probed using antibodies to the indicated proteins.
**B.** Western blot of whole-cell lysates extracted from human vein endothelial cells (HUVEC) expressing lentivirally supplied nontargeting shRNA (NT), Shoc2 shRNA (KD), Shoc2 shRNA together with Shoc2-tagRFP (SE), or Shoc2 shRNA together with Shoc2(S2G)-tagRFP (S2G) for 3 days. Cell lysates were probed using antibodies to the indicated proteins.
The results in each panel are representative of at least three independent experiments.
**C.** Western blot of whole-cell lysates extracted from human fibroblasts expressing lentivirally supplied nontargeting shRNA (NT), Shoc2 shRNA (KD), Shoc2 shRNA together with Shoc2-tagRFP (SE), or Shoc2 shRNA together with Shoc2(S2G)-tagRFP (S2G) for 3 days. Cell lysates were analyzed using the indicated antibodies.

**Figure S8. Transcriptome analysis of data sets in Figure 8.**
**A.** Gene set enrichment analysis (GSEA) presented as a score plot with NES showing upregulation of selected genes in the "Oxidative Phosphorylation" signature of Shoc2-depleted HDLEC (Shoc2 KD) compared with control (non-targeting, NT). The total height of the curve indicates the extent of enrichment (ES), with the normalized enrichment score (NES), the false discovery rate (FDR), and the p-value. Source data are provided as a source data file.
**B.** Heatmap showing differentially expressed "Oxidative Phosphorylation" genes in HDLEC expressing non-targeting shRNA, Shoc2 shRNA, or Shoc2-tRFP (p-value $\leq 0.05$, [Log2fold change] $\geq 0.0$).
Red and blue indicate increased and decreased expression for all heatmaps, respectively.

**Figure S9. Proposed model for Shoc2 in regulating LEC senescence.**
Schematic showing the working model depicting what is currently understood for the mechanisms by which Shoc2 modulates LEC homeostasis and senescence. Shoc2 prevents excessive activation of the mTORC1 complex. Shoc2 loss leads to increased Raptor binding to mTOR, increased mTORC1 signaling, mitochondrial fission, and potential leak of mitochondrial double-stranded RNA. As a result, mt-dsRNAs trigger an innate immune response, initiating sterile inflammation and cell

senescence. Our model highlights a critical role for Shoc2 in regulating cell signaling, mitochondrial dynamics, and cell senescence.

**Supplementary Video 1. Time-lapse imaging of normal secondary sprouts in a wild-type sibling zebrafish larva.**

Mid-trunk of a wild-type sibling *Tg(mrc1a:eGFP)*$^{y251}$, *Tg(kdrl:mCherry)*$^{y171}$ embryo was imagined during 32-50 hpf.

**Supplementary Video 2. Time-lapse imaging of defective secondary sprouts in a *shoc2*$^{SA/SA}$ mutant zebrafish larva.**

Mid-trunk of a *shoc2*$^{SA/SA}$ mutant *Tg(mrc1a:eGFP)*$^{y251}$, *Tg(kdrl:mCherry)*$^{y171}$ embryo was imagined during 32-50 hpf.

**Supplementary Video 3. Time-lapse imaging of normal primary sprouts in a wild-type sibling zebrafish larva.**

Mid-trunk of a wild-type sibling *Tg(fli1a:eGFP)*$^{y1}$ embryo was imagined during 24-36 hpf.

**Supplementary Video 4. Time-lapse imaging of normal primary sprouts in a *shoc2*$^{SA/SA}$ mutant zebrafish larva.**
Mid-trunk of a *shoc2*$^{SA/SA}$ mutant *Tg(fli1a:eGFP)*$^{y1}$ embryo was imagined during 24-36 hpf.

**Table S1**

| Antibody | Manufacturer | Cat# |
|---|---|---|
| Phospho-ERK1/2 (Thr202/Tyr204) | Proteintech, Rosemont, IL | 28733-1-AP |
| ERK1/2 | Proteintech, Rosemont, IL | 11257-1-AP |
| GAPDH | Proteintech, Rosemont, IL | 60004-1-1g |
| Shoc2 (Sur-8) (e-4) | SCBT, Santa Cruz, CA | SC-514888 |
| β-Actin (C4) | SCBT, Santa Cruz, CA | SC-47778 |
| mTOR mouse ( used for IP) | Proteintech, Rosemont, IL | 66888-1-1g |
| mTOR rabbit | Proteintech, Rosemont, IL | 28273-1-AP |
| Phosphor-mTOR (Ser2448) | Proteintech, Rosemont, IL | 67778-1-1g |
| Raptor | Proteintech, Rosemont, IL | 20984-1-AP |
| Rictor | CST, Danvers, MA | 2114 |
| phospho-p70S6K (Thr389) | Proteintech, Rosemont, IL | 28735-1-AP |
| p70S6K | Proteintech, Rosemont, IL | 14485-1-AP |
| Phospho-4E-BP1 (Thr37) | Proteintech, Rosemont, IL | 81812-4-RR |
| 4E-BP1 | Proteintech, Rosemont, IL | 60246-1-1g |
| STAT | Proteintech, Rosemont, IL | 10144-2-AP |
| Phospho-STAT1 (Tyr701) (58D6) | CST, Danvers, MA | 9167S |
| JAK1 | CST, Danvers, MA | 3344T |
| phospho-JAK1 (Tyr1034/1035) (D7N4Z) | CST, Danvers, MA | 74129T |
| IRF1 | Proteintech, Rosemont, IL | 11335-1-AP |
| IRF7 | Proteintech, Rosemont, IL | 22392-1-AP |
| phospho-IRF7 (Ser477) | CST, Danvers, MA | 12390T |
| IRF9 | Proteintech, Rosemont, IL | 14167-1-AP |
| MDA5 | Proteintech, Rosemont, IL | 21775-1-AP |
| MAVS | Proteintech, Rosemont, IL | 14341-1-AP |
| Phospho-MDA5 (Ser388) | Gift from Dr. Michaela Guck | |
| BST2 | Proteintech, Rosemont, IL | 13560-1-AP |
| RSAD2 | Proteintech, Rosemont, IL | 28089-1-AP |
| MX1 | Proteintech, Rosemont, IL | 13750-1-AP |
| IFN-gamma | Proteintech, Rosemont, IL | 15365-1-AP |
| ISG15 | Proteintech, Rosemont, IL | 15981-1-AP |
| USP18 | Avivasystemsbio, San Diego, CA | ARP46253-P050 |
| IFIT1 | Proteintech, Rosemont, IL | 23247-1-AP |
| HERC5 | Proteintech, Rosemont, IL | 22612-1-AP |
| Drp1 | CST, Danvers, MA | 8570 |
| phospho-Drp1 (Ser616) | SAB, Greenbelt, MD | 12749 |
| Opa1 | Proteintech, Rosemont, IL | 27733-1-AP |
| MFN1 | Proteintech, Rosemont, IL | 13798-1-AP |
| MFF | Proteintech, Rosemont, IL | 17090-1-AP |

| | | |
|---|---|---|
| p21 | Proteintech, Rosemont, IL | 10355-1-AP |
| p53 | Proteintech, Rosemont, IL | 10442-1-AP |
| Cleaved caspase 3 | Proteintech, Rosemont, IL | 25128-1-AP |
| phospho-p53 | Proteintech, Rosemont, IL | 28961-1-AP |
| Peroxidase-conjugated AffiniPure F(ab')2 Fragment Goat Anti-Rabbit and -Mouse IgG (H+L) | Jackson ImmunoResearch, West Grove, PA | 111-035-006 (rabbit) 115-035-006 (mouse) |

**Table S2**

| Gene | Forward Primer (5' to 3') | Reverse Primer (5' to 3') | Accession No./REF |
|---|---|---|---|
| BST2 | GTGGAGCGACTGAGAAGAGAAA | CGGACCTTCCAAGATGTGCC | NM_004335.4 |
| CXCL10 | AGCAGTTAGCAAGGAAAGGTCT | GGAGGATGGCAGTGGAAGTC | NM_001565.4 |
| DDX60 | GGTGTTTTCACCAGGGAGTATCG | CCAGTTTTGGCGATGAGGAGCA | NM_017631.6 |
| HERC6 | TGTGGACACTACCACTCCCT | CCATGGCTGTTCTTTCCCCA | NM_017912.4 |
| IFI27 | ATCAGCAGTGACCAGTGTGG | TGGCCACAACTCCTCCAATC | NM_001366993.1 |
| IFI6 | AAGGCGGTATCGCTTTTCTTG | TTTTCTTACCTGCATCCTTACCC | NM_022873.3 |
| IFIH1 (MDA5) | GCTGAAGTAGGAGTCAAAGCCC | CCACTGTGGTAGCGATAAGCAG | NM_022168.4 |
| IRF7 | TACCATCTACCTGGGCTTCG | AGGGTTCCAGCTTCACCA | NM_001572.5 |
| IRF9 | GCATCAGGCAGGGCACGCTGCACC | GCCTGCATGTTTCCAGGGAATCCG | NM_001385400.1 |
| ISG15 | TGGCGGGCAACGAATT | GGGTGATCTGCGCCTTCA | NM_005101.4 |
| MX1 | ACCTGTGCAGCCAGTATGAGG | AGCCCGCAGGGAGTCAA | NM_001144925.2 |
| DDX-58 (RIG-I) | CACCTCAGTTGCTGATGAAGGC | GTCAGAAGGAAGCACTTGCTACC | NM_014314.4 |
| TRIM14 | CAACAGGGTCTGGAGTATCAGC | CTTGACGGGCTCAAAAGAGAGG | NM_014788.4 |
| TRIM25 | AAAGCCACCAGCTCACATCCGA | GCGGTGTTGTAGTCCAGGATGA | NM_005082.5 |
| USP18 | CAGACCCTGACAATCCACCT | AGCTCATACTGCCCTCCAGA | NM_017414.4 |
| SHOC2 | TCAGTGGTGTATAGGCTGGATTCT | GCTACATCCAGCGTAATGAGGT | Geng and Gao, 2019 |
| PROX1A | GCTCCAATATGCTGAAGACC | ATCGTTGATGGCTTGACGTG | Park, H.J, 2018 |
| LYVE1 | GCAAGGACCAAGTTGAAACAGCCTTG | CTGGAATGCACGAGTTAGTCCAAGTA | Park, H.J, 2018 |
| GAPDH | GGTGGTCTCCTCTGACTTCA | GTTGCTGTAGCCAAATTCGT | Kim et al, 2017 |

**Table S3**

| Gene | Forward Primer (5' to 3') | Reverse Primer (5' to 3') | Accession No. |
|---|---|---|---|
| Shoc2 Δ22 | CCATCAAGGAGCTGACCCAG | AGTCAGGTAGGCTGGTCAGA | NM_00104478.1 |
| Shoc2 SA/SA | TCCCTTTTGGCATTTTCTCTCG | GAGTTTGTTCAGCCAGCATCC | NM_00104478.1 |
| Shoc2 SA/SA Sequencing | CGGTTTTACCAGAGGGGCTT | CCACCATACTGGTCCACGTC | NM_00104478.1 |

**Table S4**

| Gene | SENSE (5' to 3') | ANTISENSE (5' to 3') | Reference |
|---|---|---|---|
| MDA5 | GUUAUAGUUCUUGUCAAUAUU | UAUUGACAAGAACUAUAACUU | Kuo, RL, 2013 PlosOne |
| MAVSa | CCACCUUGAUGCCUGUGAAUU | UUCACAGGCAUCAAGGUGGUU | Cheng G, 2007, PNAS |
| MAVSb | CAGAGGAGAAUGAGUAUAAUU | UUAUACUCAUUCUCCUCUGUU | Seth, R.B. 2005, Cell |

Figure S1

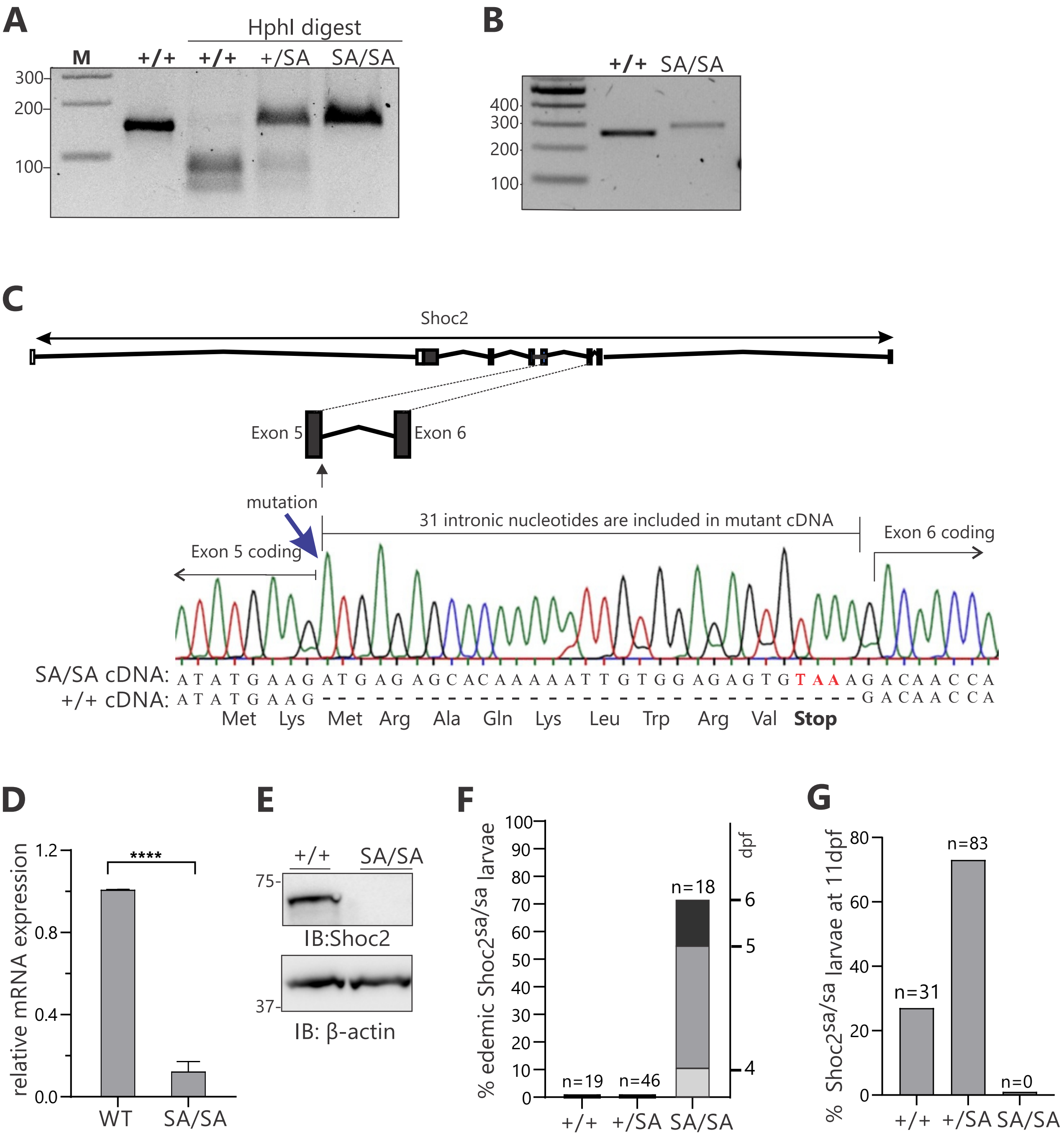
A
HphI digest
M
+/+
+/+
+/SA
SA/SA
300
200
100
B
+/+
SA/SA
400
300
200
100
C
Shoc2
Exon 5
Exon 6
mutation
31 intronic nucleotides are included in mutant cDNA
Exon 5 coding
Exon 6 coding
SA/SA cDNA: A T A T G A A G A T G A G A G C A C A A A A A T T G T G G A G A G T G T A A A G A C A A C C A
+/+ cDNA: A T A T G A A G - - - - - - - - - - - - - - - - - - - - - - - - - - - - - - - G A C A A C C A
Met Lys Met Arg Ala Gln Lys Leu Trp Arg Val Stop
D
relative mRNA expression
1.2
0.8
0.4
0.0
****
WT
SA/SA
E
+/+
SA/SA
75
IB:Shoc2
37
IB: β-actin
F
% edemic Shoc2sa/sa larvae
100
90
80
70
60
50
40
30
20
10
0
n=19
n=46
n=18
dpf
6
5
4
+/+
+/SA
SA/SA
G
% Shoc2sa/sa larvae at 11dpf
80
60
40
20
0
n=31
n=83
n=0
+/+
+/SA
SA/SA

### Figure S2

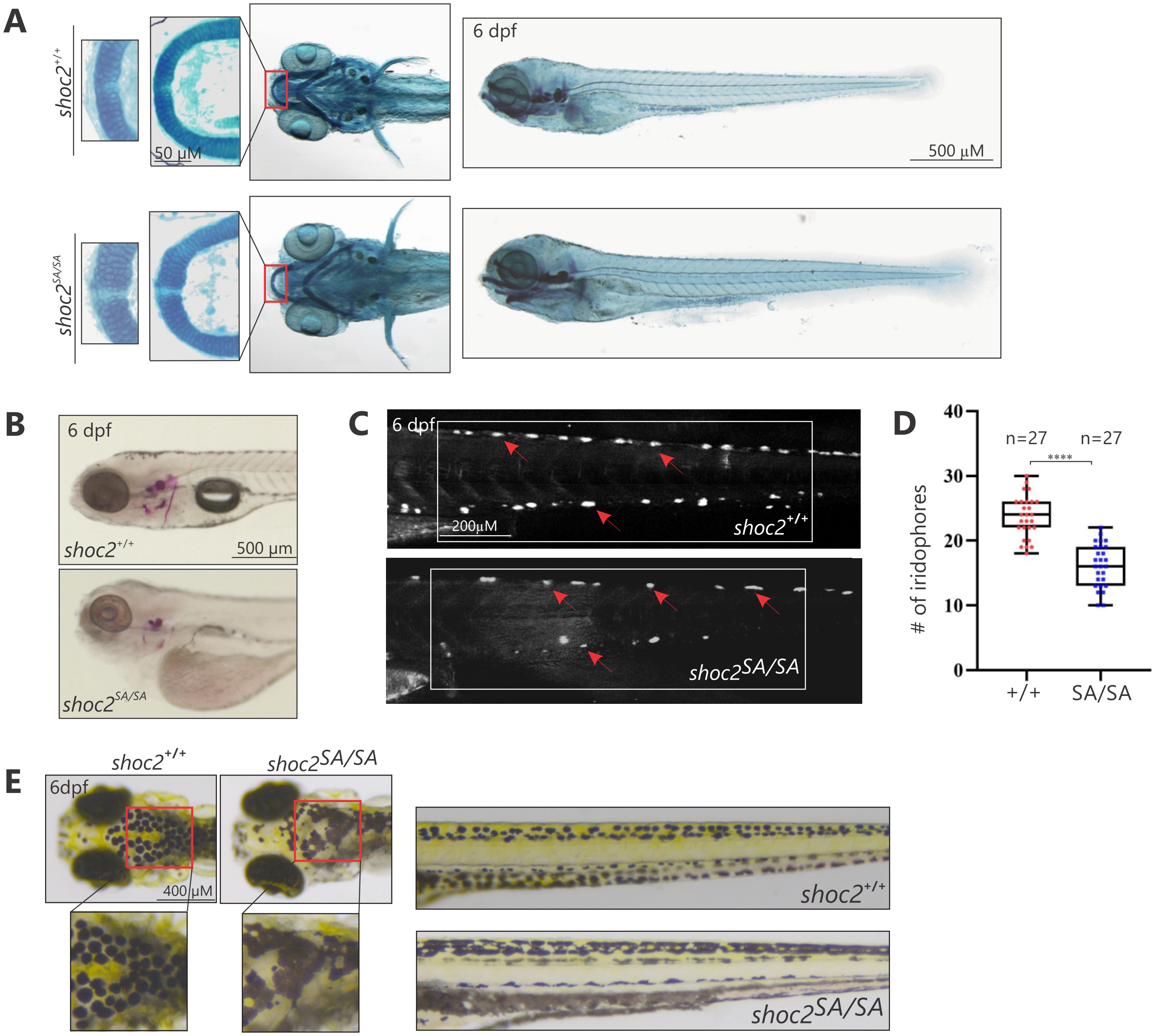
A
shoc2+/+
50 µM
6 dpf
500 µM
shoc2SA/SA
B
6 dpf
shoc2+/+
500 µm
shoc2SA/SA
C
6 dpf
200µM
shoc2+/+
shoc2SA/SA
D
n=27
n=27
****
### of iridophores
40
30
20
10
0
+/+
SA/SA
E
shoc2+/+
shoc2SA/SA
6dpf
400 µM
shoc2+/+
shoc2SA/SA

Figure S3

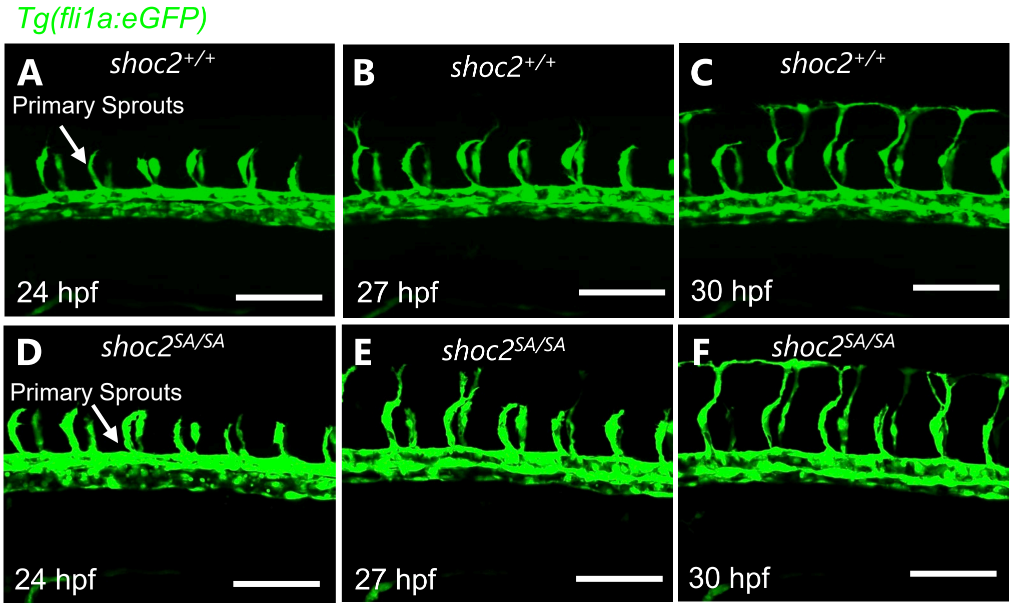

Tg(fli1a:eGFP)
A
shoc2+/+
Primary Sprouts
24 hpf
B
shoc2+/+
27 hpf
C
shoc2+/+
30 hpf
D
shoc2SA/SA
Primary Sprouts
24 hpf
E
shoc2SA/SA
27 hpf
F
shoc2SA/SA
30 hpf

Figure S4

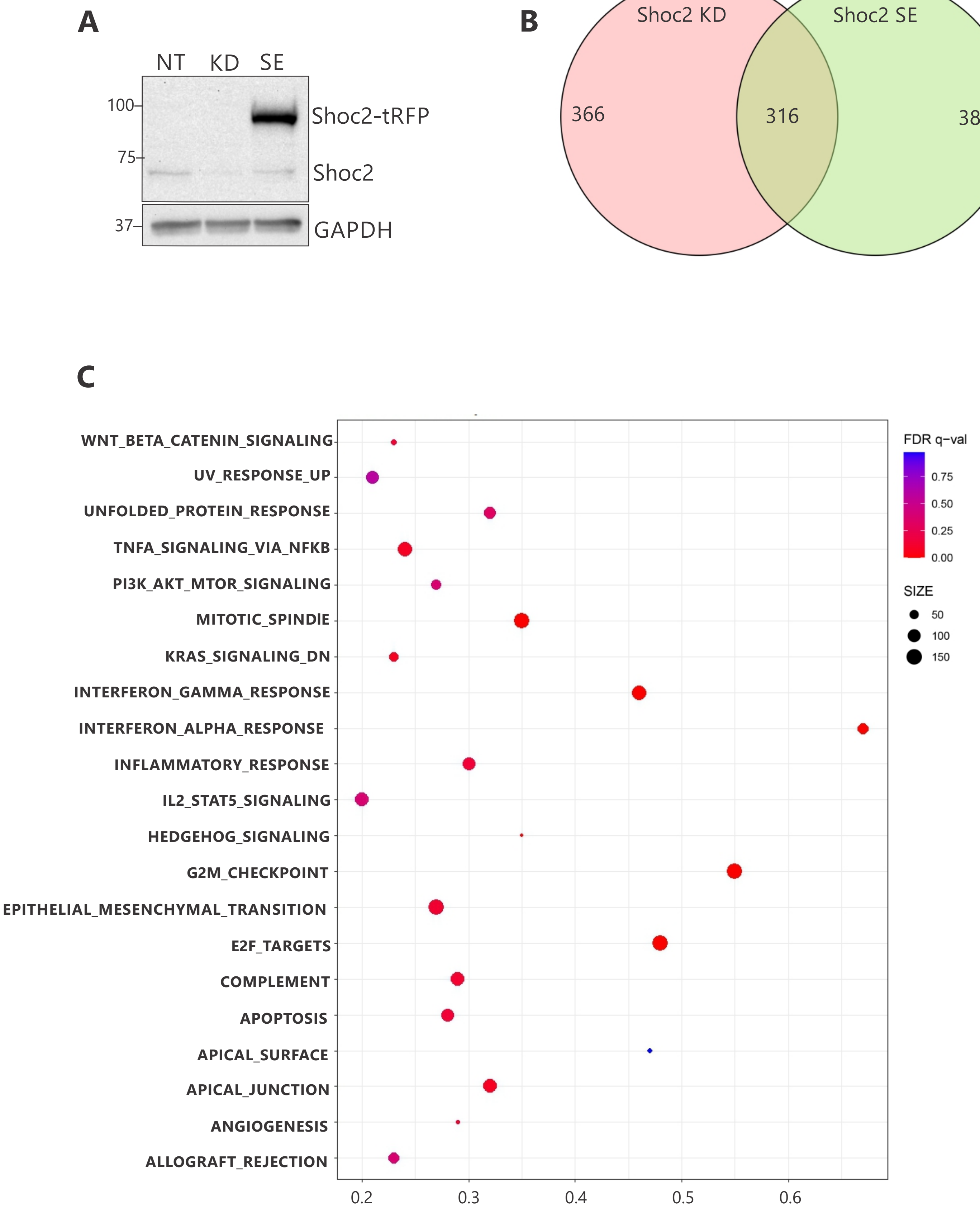
A
NT
KD
SE
100
75
37
Shoc2-tRFP
Shoc2
GAPDH
B
Shoc2 KD
Shoc2 SE
366
316
383
C
WNT_BETA_CATENIN_SIGNALING
UV_RESPONSE_UP
UNFOLDED_PROTEIN_RESPONSE
TNFA_SIGNALING_VIA_NFKB
PI3K_AKT_MTOR_SIGNALING
MITOTIC_SPINDIE
KRAS_SIGNALING_DN
INTERFERON_GAMMA_RESPONSE
INTERFERON_ALPHA_RESPONSE
INFLAMMATORY_RESPONSE
IL2_STAT5_SIGNALING
HEDGEHOG_SIGNALING
G2M_CHECKPOINT
EPITHELIAL_MESENCHYMAL_TRANSITION
E2F_TARGETS
COMPLEMENT
APOPTOSIS
APICAL_SURFACE
APICAL_JUNCTION
ANGIOGENESIS
ALLOGRAFT_REJECTION
0.2
0.3
0.4
0.5
0.6
FDR q−val
0.75
0.50
0.25
0.00
SIZE
50
100
150

### Figure S5

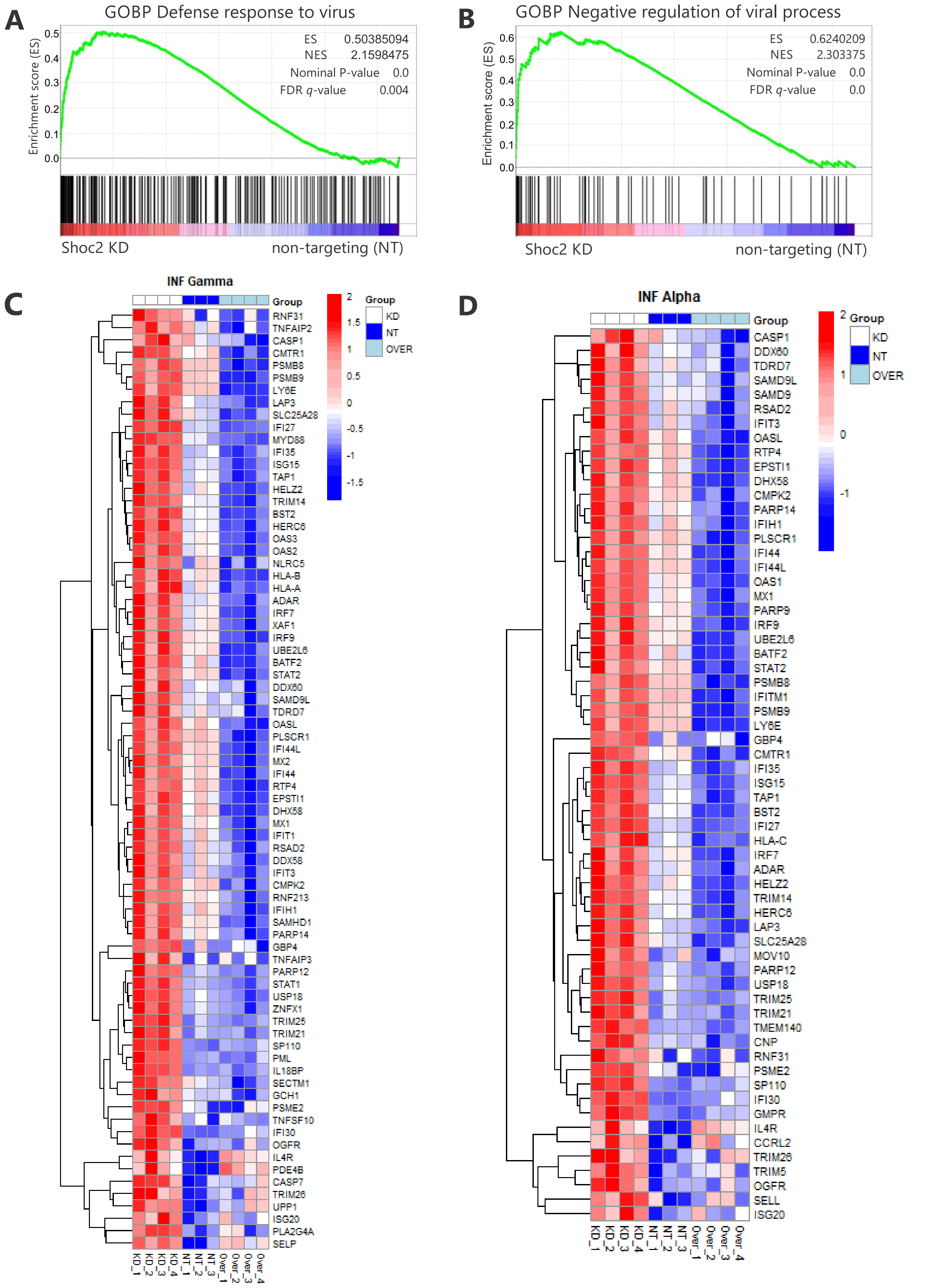

A
GOBP Defense response to virus
ES 0.50385094
NES 2.1598475
Nominal P-value 0.0
FDR q-value 0.004
Enrichment score (ES)
0.5
0.4
0.3
0.2
0.1
0.0
Shoc2 KD
non-targeting (NT)
B
GOBP Negative regulation of viral process
ES 0.6240209
NES 2.303375
Nominal P-value 0.0
FDR q-value 0.0
Enrichment score (ES)
0.6
0.5
0.4
0.3
0.2
0.1
0.0
Shoc2 KD
non-targeting (NT)
C
INF Gamma
Group
KD
NT
OVER
2
1.5
1
0.5
0
-0.5
-1
-1.5
RNF31
TNFAIP2
CASP1
CMTR1
PSMB8
PSMB9
LY6E
LAP3
SLC25A28
IFI27
MYD88
IFI35
ISG15
TAP1
HELZ2
TRIM14
BST2
HERC6
OAS3
OAS2
NLRC5
HLA-B
HLA-A
ADAR
IRF7
XAF1
IRF9
UBE2L6
BATF2
STAT2
DDX60
SAMD9L
TDRD7
OASL
PLSCR1
IFI44L
MX2
IFI44
RTP4
EPSTI1
DHX58
MX1
IFIT1
RSAD2
DDX58
IFIT3
CMPK2
RNF213
IFIH1
SAMHD1
PARP14
GBP4
TNFAIP3
PARP12
STAT1
USP18
ZNFX1
TRIM25
TRIM21
SP110
PML
IL18BP
SECTM1
GCH1
PSME2
TNFSF10
IFI30
OGFR
IL4R
PDE4B
CASP7
TRIM26
UPP1
ISG20
PLA2G4A
SELP
KD_1
KD_2
KD_3
KD_4
NT_1
NT_2
NT_3
Over_1
Over_2
Over_3
Over_4
D
INF Alpha
Group
KD
NT
OVER
2
1
0
-1
CASP1
DDX60
TDRD7
SAMD9L
SAMD9
RSAD2
IFIT3
OASL
RTP4
EPSTI1
DHX58
CMPK2
PARP14
IFIH1
PLSCR1
IFI44
IFI44L
OAS1
MX1
PARP9
IRF9
UBE2L6
BATF2
STAT2
PSMB8
IFITM1
PSMB9
LY6E
GBP4
CMTR1
IFI35
ISG15
TAP1
BST2
IFI27
HLA-C
IRF7
ADAR
HELZ2
TRIM14
HERC6
LAP3
SLC25A28
MOV10
PARP12
USP18
TRIM25
TRIM21
TMEM140
CNP
RNF31
PSME2
SP110
IFI30
GMPR
IL4R
CCRL2
TRIM26
TRIM5
OGFR
SELL
ISG20
KD_1
KD_2
KD_3
KD_4
NT_1
NT_2
NT_3
Over_1
Over_2
Over_3
Over_4

### Figure S6

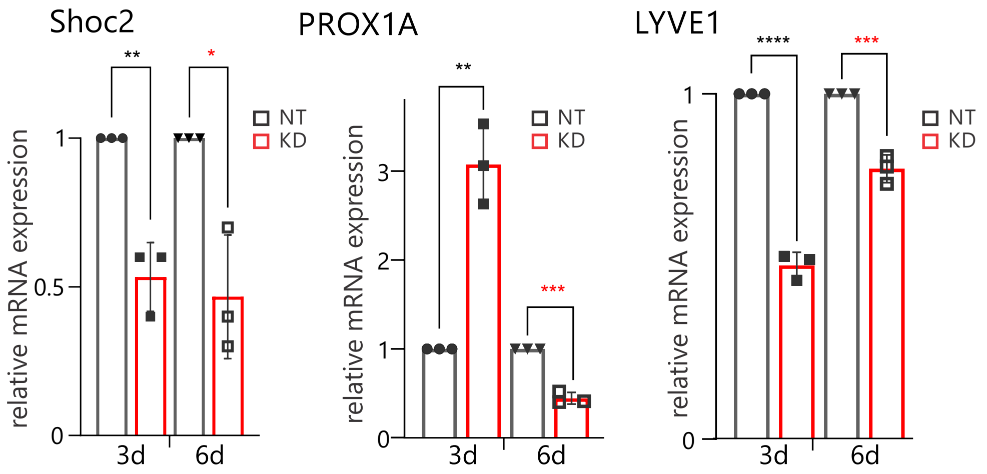
Shoc2
PROX1A
LYVE1
relative mRNA expression
NT
KD
3d
6d
0
0.5
1
2
3
**
*
***
****

Figure S7

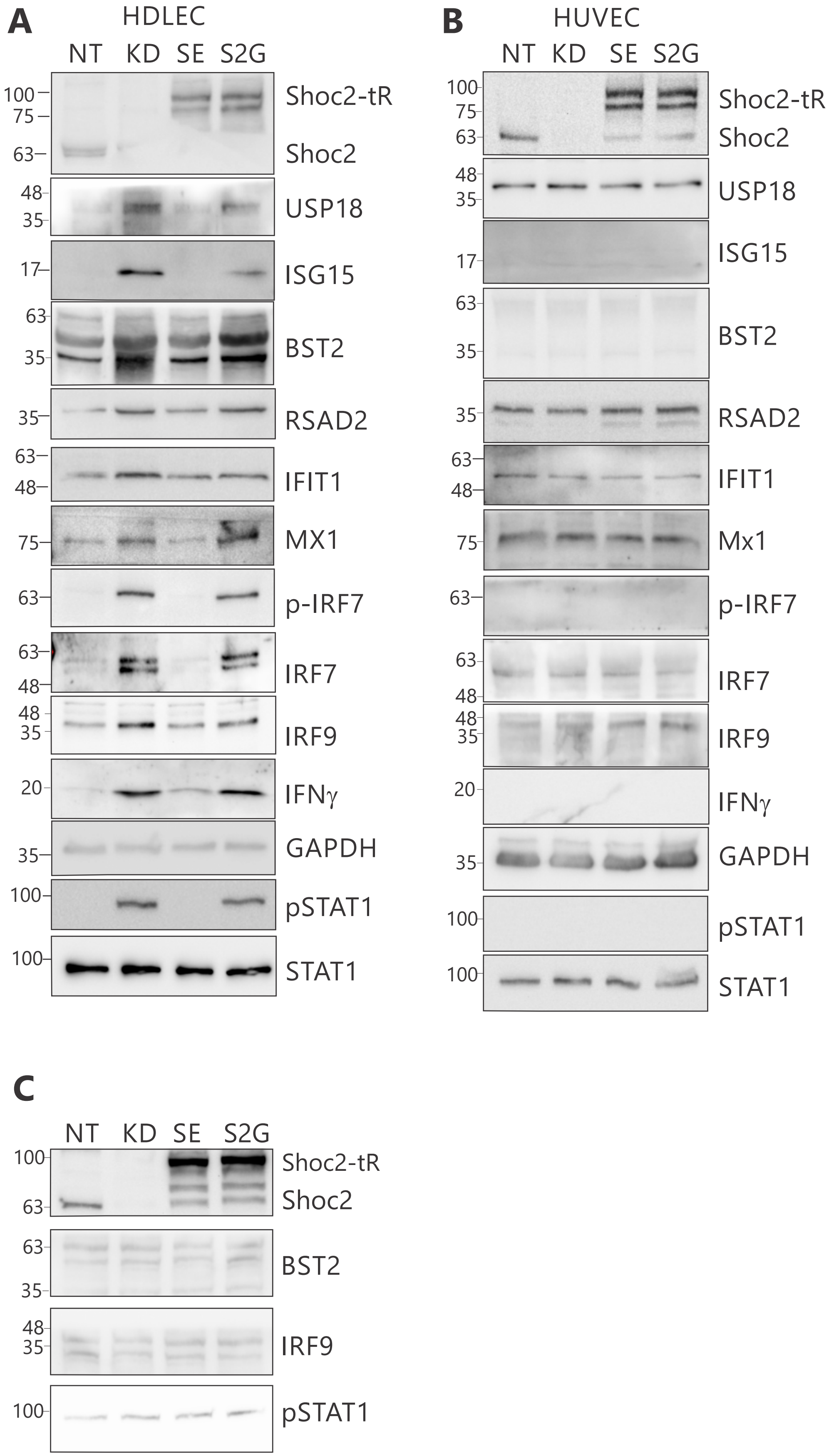
A
HDLEC
NT KD SE S2G
100
75
63
Shoc2-tR
Shoc2
48
35
USP18
17
ISG15
63
35
BST2
35
RSAD2
63
48
IFIT1
75
MX1
63
p-IRF7
63
48
IRF7
48
35
IRF9
20
IFNγ
35
GAPDH
100
pSTAT1
100
STAT1
B
HUVEC
NT KD SE S2G
100
75
63
Shoc2-tR
Shoc2
48
35
USP18
17
ISG15
63
35
BST2
35
RSAD2
63
48
IFIT1
75
Mx1
63
p-IRF7
63
48
IRF7
48
35
IRF9
20
IFNγ
35
GAPDH
100
pSTAT1
100
STAT1
C
NT KD SE S2G
100
63
Shoc2-tR
Shoc2
63
35
BST2
48
35
IRF9
100
pSTAT1

### Figures S8

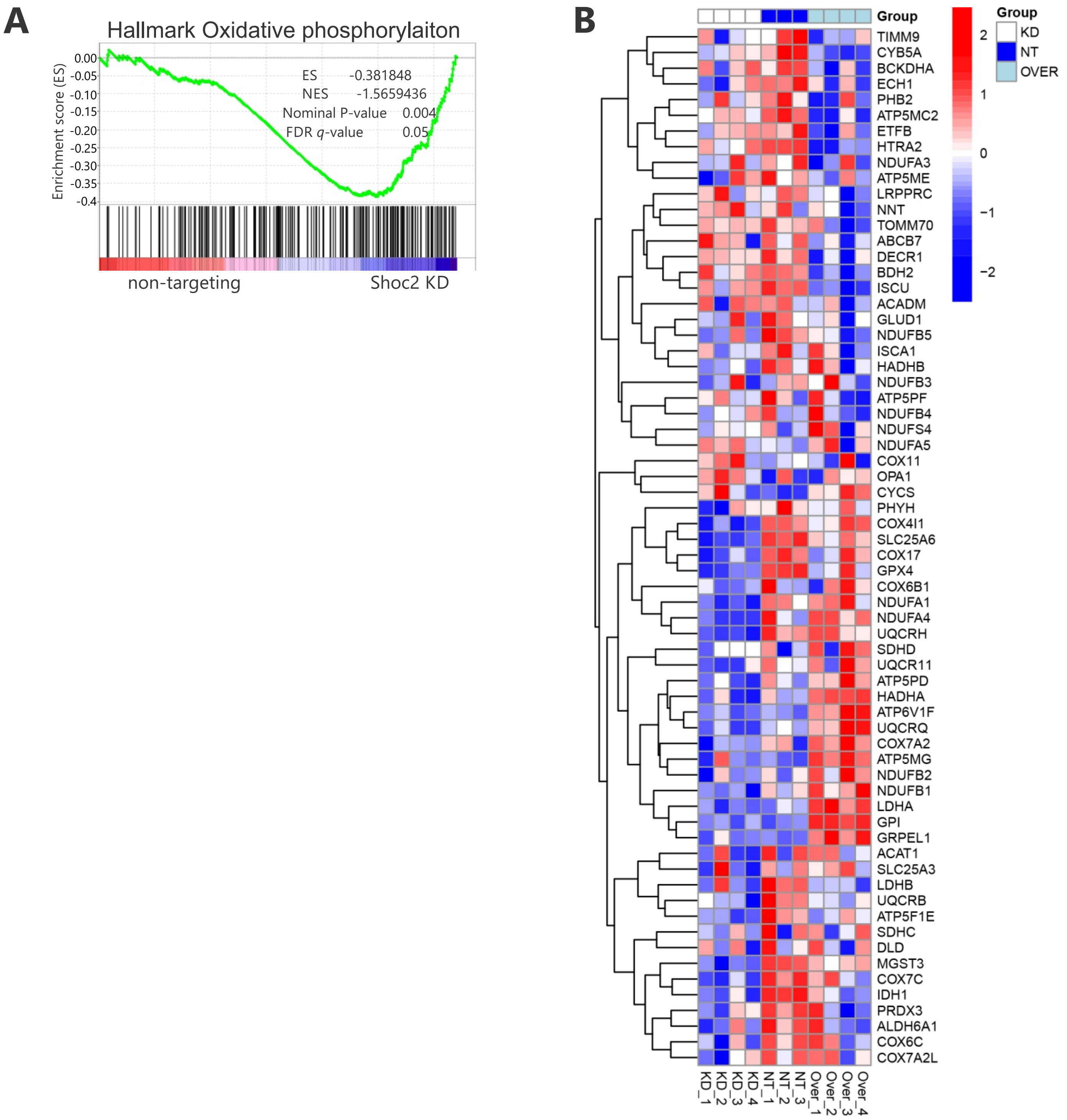
A
Hallmark Oxidative phosphorylaiton
Enrichment score (ES)
ES -0.381848
NES -1.5659436
Nominal P-value 0.004
FDR q-value 0.05
non-targeting
Shoc2 KD
B
Group
KD
NT
OVER
TIMM9
CYB5A
BCKDHA
ECH1
PHB2
ATP5MC2
ETFB
HTRA2
NDUFA3
ATP5ME
LRPPRC
NNT
TOMM70
ABCB7
DECR1
BDH2
ISCU
ACADM
GLUD1
NDUFB5
ISCA1
HADHB
NDUFB3
ATP5PF
NDUFB4
NDUFS4
NDUFA5
COX11
OPA1
CYCS
PHYH
COX4I1
SLC25A6
COX17
GPX4
COX6B1
NDUFA1
NDUFA4
UQCRH
SDHD
UQCR11
ATP5PD
HADHA
ATP6V1F
UQCRQ
COX7A2
ATP5MG
NDUFB2
NDUFB1
LDHA
GPI
GRPEL1
ACAT1
SLC25A3
LDHB
UQCRB
ATP5F1E
SDHC
DLD
MGST3
COX7C
IDH1
PRDX3
ALDH6A1
COX6C
COX7A2L
KD_1
KD_2
KD_3
KD_4
NT_1
NT_2
NT_3
Over_1
Over_2
Over_3
Over_4

### Figure S9

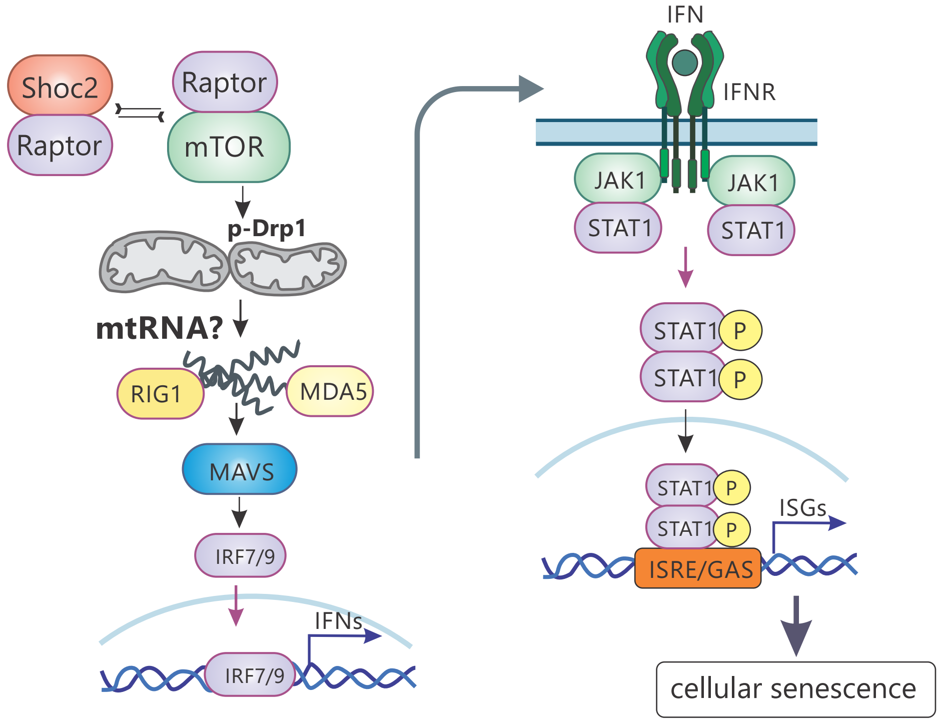
Shoc2
Raptor
Raptor
mTOR
p-Drp1
mtRNA?
RIG1
MDA5
MAVS
IRF7/9
IFNs
IRF7/9
IFN
IFNR
JAK1
JAK1
STAT1
STAT1
STAT1
P
STAT1
P
STAT1
P
STAT1
P
ISGs
ISRE/GAS
cellular senescence
